# TransfoRNA: Navigating the Uncertainties of Small RNA Annotation with an Adaptive Machine Learning Strategy

**DOI:** 10.1101/2024.06.19.599329

**Authors:** Yasser Taha, Julia Jehn, Mustafa Kahraman, Maurice Frank, Marco Heuvelman, Rastislav Horos, Christopher Yau, Bruno Steinkraus, Tobias Sikosek

## Abstract

Small RNAs hold crucial biological information and have immense diagnostic and therapeutic value. While many established annotation tools focus on microRNAs, there are myriads of other small RNAs that are currently underutilized. These small RNAs can be difficult to annotate, as ground truth is limited and well-established mapping and mismatch rules are lacking.

TransfoRNA is a machine learning framework based on Transformers that explores an alternative strategy. It uses common annotation tools to generate a small seed of high-confidence training labels, while then expanding upon those labels iteratively. TransfoRNA learns sequence-specific representations of all RNAs to construct a similarity network which can be interrogated as new RNAs are annotated, allowing to rank RNAs based on their familiarity. While models can be flexibly trained on any RNA dataset, we here present a version trained on TCGA (The Cancer Genome Atlas) small RNA sequences and demonstrate its ability to add annotation confidence to an unrelated dataset, where 21% of previously unannotated RNAs could be annotated. Relative to its training data, TransfoRNA could boost high-confidence annotations in TCGA by ∼50% while providing transparent explanations even for low-confidence ones. It could learn to annotate 97% of isomiRs from just single examples and confidently identify new members of other familiar classes with high accuracy, while reliably rejecting false RNAs.

All source code is available at https://github.com/gitHBDX/TransfoRNA and can be executed at Code Ocean (https://codeocean.com/capsule/5415298/). An interactive website is available at www.transforna.com.

**GRAPHICAL ABSTRACT:** 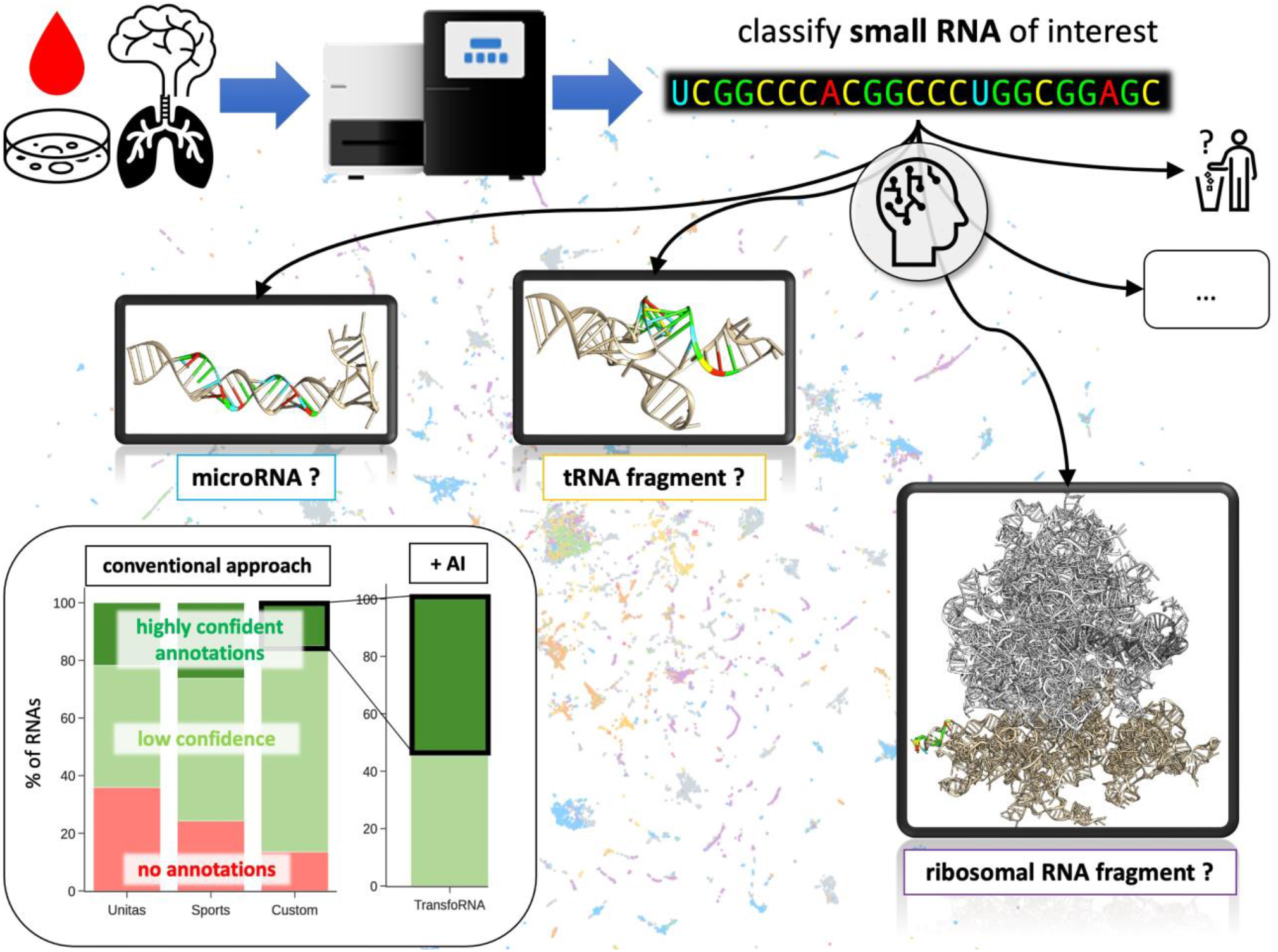

## INTRODUCTION

Small-RNA sequencing is a technology that allows for the simultaneous readout of millions of RNA molecules from biological (e.g., patient) samples for basic research as well as diagnostic or therapeutic purposes. Small RNAs are typically around 20 nucleotides (nt) long. The largest and most important group of small RNAs are microRNAs (miRNA) which are responsible for intracellular regulation and signaling (1). Other small RNAs, however, are fragments of much larger RNAs. For example, tRNAs (transfer RNAs) and rRNAs (ribosomal RNAs) are integral parts of the core cellular machinery that translates genetic information from DNA into proteins. These larger RNAs are routinely broken down into smaller fragments at the end of their life cycle. Some research suggests that some of these RNA fragments have a function of their own (2–9). Other classes of RNAs are also found in lower quantities (10–12).

The traditional focus of small-RNA sequencing (also: next-generation sequencing, NGS, or RNA-seq) has been the mapping to known miRNA families, while discarding the reads of most other RNA classes. It is also common to aggregate multiple miRNA sequence isoforms (isomiRs) that map to the same reference miRNA. However, this will obscure any differential expression between isomiRs summed up under the same name. Therefore, there is an underutilized opportunity to treat each individual small RNA as a potential regulator or marker of important biology or disease (8, 13–17).

Several published software tools exist for such conventional, knowledge-based annotation (KBA). KBA approaches make use of expert-curated databases such as miRbase (18) and map against a human reference genome (19, 20). In **Figure 1** we compare two of these (Sports(19) and Unitas(21)) with the annotation approach used herein for TransfoRNA (labeled “custom”), demonstrating that for the most part they are similar in how many RNAs are annotated as one of several major classes. The user needs to specify the tolerated number of mismatches when mapping against the reference databases. At the strictest settings (no mismatch), more than 70% of RNAs would not be annotated. At default settings, that number can still exceed 30%. RNAs that fail to be annotated may be novel in a biological sense but could also be technical artefacts of the NGS technology (22). Removal of these artefacts, however, remains a challenge towards full use of small-RNA datasets. Additional challenges exist. Due to the short length of these RNAs there is a risk of false positive mapping hits and the possibility of mismatches, i.e. due to patient-specific genomic mutations, post-transcriptional modifications, as well as simple sequencing (base-calling) mistakes. Devising a rule-based algorithm attempting to capture all such possibilities and weighing their relative probabilities is a daunting task. Therefore, there is a clear need for an improved annotation strategy.

**Figure 1:**
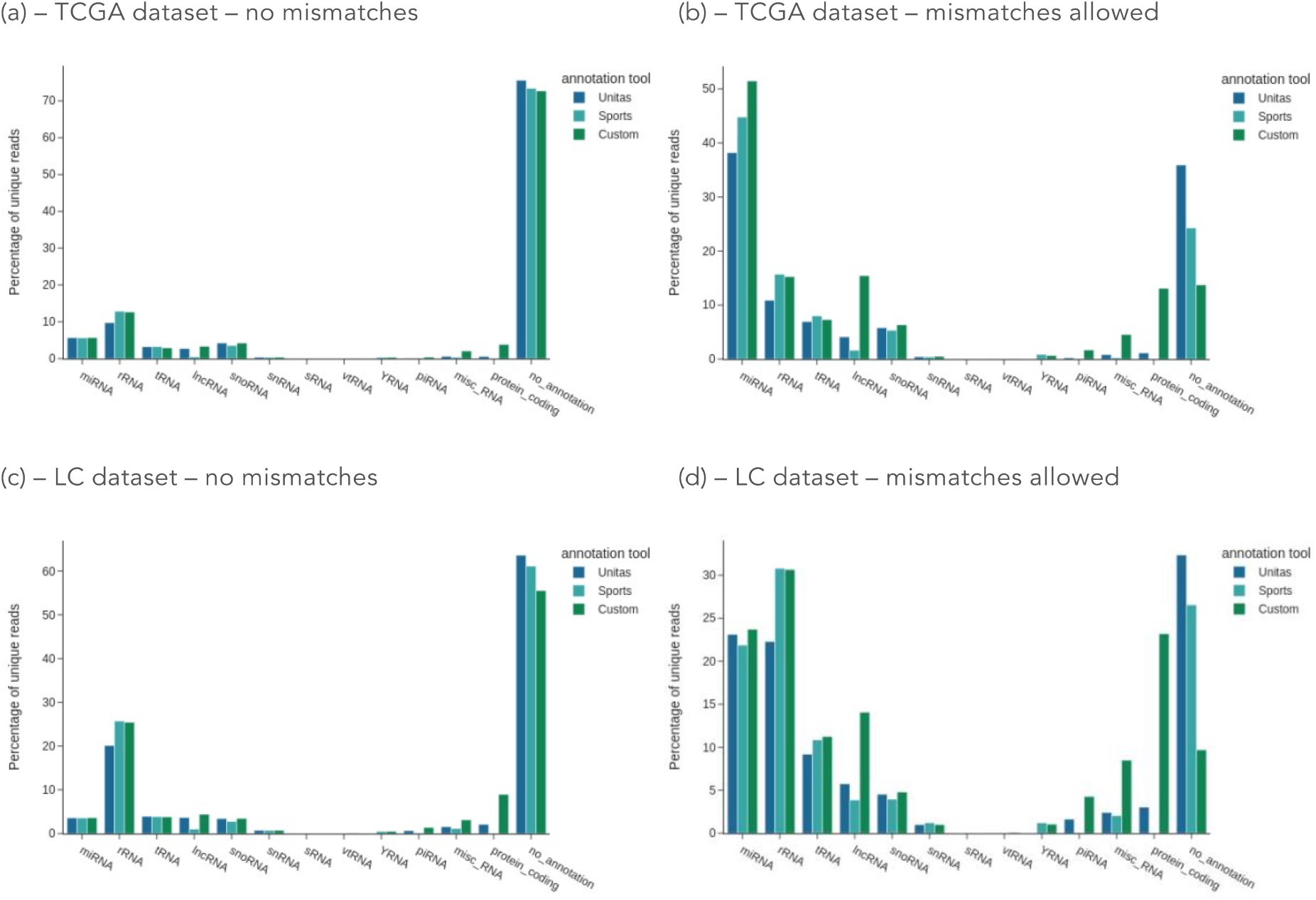
Knowledge-based annotation (KBA) tools require the choice of how many mismatches to tolerate during mapping to the transcriptomic reference. The strictest settings (a+c) require perfect matches and are used here to define “high-confidence” (HICO) annotations, which are used for training TransfoRNA. At default settings (b+d), most tools allow mismatches to increase the number of annotated RNAs. These additional annotations were considered “low confidence” (LOCO) here and together with the “no annotation” set were (re-)annotated by TransfoRNA. TCGA: the cancer genome atlas dataset. LC: lung cancer dataset. “Custom” refers to the KBA method developed here for the purpose of training TransfoRNA machine learning models.

Machine learning (ML), a subfield of artificial intelligence (AI), offers a flexible solution for decision making. A machine learning model could learn general RNA sequence patterns from high-confidence biological ground truth annotations and then apply those to the annotation of any other RNA sequence. Ideally such an approach would allow the models to also recognize any non-biological RNAs (artefacts) as particularly unusual or novel. Another advantage of machine learning is the ease of incorporating additional potentially relevant information into the learning process, such as RNA secondary structure.

However, the main limitation for machine learning is the availability of correct training labels (ground truth) and therefore machine learning should not be seen as a replacement for knowledge-based annotation methods, but as an extension. Reports of potentially mis-labeled database entries, which have been pointed out especially for miRNAs (23–28), are further justification for a conservative approach towards annotation ground truth. Another important consideration is the open-endedness of the annotation task. The trained models are likely to at some point encounter classes of RNA that they have not been trained on. Instead of randomly predicting one of the known classes, an ML model should ideally let the user know when it deems an RNA unfamiliar or novel.

Our approach combines both methodologies, expert knowledge, and cutting-edge machine learning, to extrapolate patterns from a small number of high-confidence labels and transfer those annotations to RNA sequences that conventional methods are not able to reliably annotate. Exploiting similarities between biological sequences and natural language, we found that the Transformer deep-learning architecture (29) works well for classification on small-RNA sequences. Transformers have been a key technology behind recent advances in large language models (LLMs), like GPT (30) or LLaMA (31). Transformers have previously already been adapted to other biological sequence data such as protein sequences (32, 33), long RNAs (34), and genomic DNA (35). We compare different formats for presenting both the small-RNA inputs, as well as the classification labels, to the models.

On the side of the user, no parameter choices such as the number of allowed genomic mismatches need to be made. Instead, the machine learning model should be trained on only the most confident annotations (no mismatches, no potential alternative origins) and will iteratively learn to confidently annotate most of a dataset.

We show that TransfoRNA models reliably identify novel RNAs from various sources (biological, technical, and computer-generated) and assist the user in making informed decisions based on the degree of novelty, the consensus among an ensemble of models, and the most similar RNAs from the training set.

TransfoRNA was trained on small RNA sequences from one dataset (TCGA tissue samples) and demonstrated excellent performance on small RNA sequences in a different dataset (blood samples from lung patients) during inference. We provide updated annotations for the TCGA dataset, which previously has only been analyzed with a focus on miRNAs (36).

## METHODS

TransfoRNA is a <u>Transfo</u>rmer-based <u>RNA</u> sequence annotation strategy for small RNAs, as they would be obtained from NGS experiments. It works in combination with conventional, knowledge-based approaches.

TransfoRNA requires small RNA sequences as input, which are taken from the large public TCGA (The Cancer Genome Atlas) (37) dataset which encompasses RNAs measured in tissue samples (healthy and cancerous) from multiple organs and many patients. A total of 75080 RNAs with a sequence length between 18 and 30 nt and detected in at least 30 samples per TCGA category were used.

Each RNA sequence was processed by a customized knowledge-based annotation (KBA) pipeline that assigns RNA class labels based on matches to reference sequences of specialized databases. We differentiate between eleven broad “major classes” (miRNA, rRNA, tRNA, snoRNA, protein coding (mRNA), lncRNA, YRNA, snRNA, piRNA, YRNA and vtRNA) and ∼2000 very detailed “sub-classes” (**Figure 2a**). See **SI Methods** and **Figure S1** for a detailed outline of the KBA approach. These KBAs are used as the training labels for later classification steps by machine learning. Sub-classes consider the type of precursor RNAs that the small RNA was derived from (e.g. miRNA precursor hairpins or mature ribosomal RNAs) as well as approximate positions along the longer precursor sequence (bins of consecutive positions). Where only very few members of a sub-class were observed, an augmentation process was employed to sample randomly chosen fragments from the corresponding bin on the precursor sequence (**Figure 2b**). Data augmentation is a common practice in modern machine learning (38, 39).

**Figure 2:**
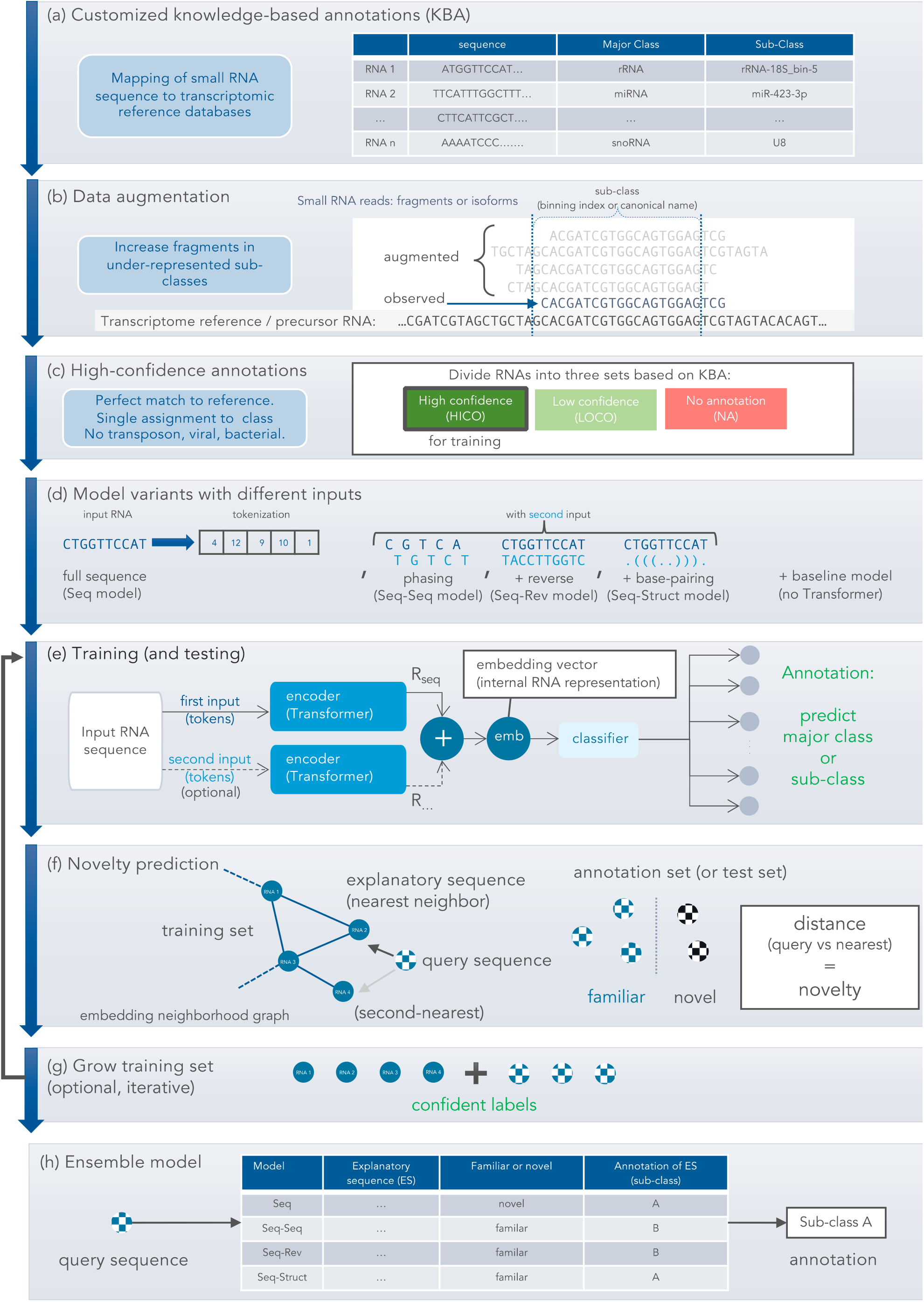
TransfoRNA annotation strategy and workflow. (a) Knowledge-based annotations (KBA) provide coarse grained major class labels and fine-grained sub-class labels within each major class. (b) Sub-classes may be defined as bins along the sequence of a much longer precursor RNA. In sub-classes where only few fragments were observed, augmentation is used to artificially produce more fragments. miRNAs are excluded from this. (c) Only high-confidence annotations are used for training, based on strict criteria. Low-confidence annotations failed these criteria. There is also a substantial number of missing annotations from KBA. (d) The RNA sequences need to be converted to numerical tokens, each representing a dinucleotide, and then are either used as input to a single Transformer module, or different representations of the same sequence are presented to two parallel Transformers. For comparison, a baseline model does not use a Transformer. (e) Each Transformer encodes the input to an embedding vector (R), two of which may be concatenated to provide the input to a classifier module, trained on major class or sub-class labels. (f) Novelty prediction assesses whether an RNA is similar to the training set and thus familiar to the model, or novel. A nearest-neighbor graph in embedding space is used to find the nearest, explanatory RNA to a given query RNA. The Normalized Levenshtein Distance (NLD) between the two is compared against a fitted threshold. (g) At this stage, the training set may be enlarged by predicted annotations that are familiar. (h) Different model variants from step d are combined to a final predictor that uses the consensus annotation among those deemed familiar.

Annotation is at its core a classification task. Machine learning classifiers need reliable ground truth labels of their input data to provide accurate predictions. To make sure that training labels are reliable, RNAs are assigned to a high-confidence (HICO) annotation set, as well as a low-confidence (LOCO) one. For the HICO set, only perfect and unique transcriptome matches were considered. In the case of miRNAs, we annotated per sub-class only the canonical reference miRNA (refmiR) from miRBase. The HICO set did not include sequence variants of these refmiRs, so-called isomiRs, as the precise criteria for considering a specific refmiR variant as a valid isomiR are still debated (40, 41). However, we have used a working definition of isomiRs similar to (42) (also see **SI Methods**) to evaluate TransfoRNA’s ability to extrapolate isomiR annotations from a single given example (see below). RNAs in the LOCO set, on the other hand, had matches with multiple reference sequences or could not be perfectly matched. Finally, one set could not be annotated even with relaxed criteria (“no annotation”, NA) (**Figure 2c**). Only HICO annotations were used for training, although the training set was allowed to grow in subsequent steps.

TransfoRNA encodes input RNA sequences (or structures) into a vector representation (i.e. embedding) that is then used to classify the sequence as an RNA class. First, each RNA sequence is encoded into a fixed-length vectorized form, which involves a tokenization step (**Figure 2d**; see **SI Methods** and **Figure S2a-I**). In analogy to natural language, each token assigns a two-letter “word” to a unique number, with the RNA alphabet only having four letters in total. In a baseline model, these tokens were encoded with a simple embedding layer followed by shallow classification (**SI Methods**). We considered the potential for alternative ways of tokenizing and encoding the sequence, e.g., splitting it into two sequences at alternating index positions (phasing), by combining the sequence with its reverse, and by pairing it with a secondary structure representation (base-pairing indicated by matching parentheses, i.e. dot-bracket notation). Each alternative input representation led to the training of a different model: baseline (plain sequence, no Transformer), Seq (plain sequence), Seq-Seq (two sequences via phasing), Seq-Rev (sequence + sequence reversed), and Seq-Struct (sequence + secondary structure). See **Figure 2d** and **Figure S2**.

The architecture of the TransfoRNA model is shown in **Figure 2e**, and in more detail in **Figure S2** RNA sequences were converted to embeddings (“emb”) by encoding a single, tokenized RNA with a Transformer module, or by feeding two alternative versions of the RNA into parallel Transformers and concatenating their output embeddings, before feeding that embedding into a multi-layer (deep), neural-network, classifier to predict the major class or sub-class of the RNA.

When using the trained models for inference, i.e., to annotate a new query RNA, the novelty of that RNA was assessed based on its embedding’s proximity to embeddings of RNAs in the training set (see below). Based on this distance and a previously fitted threshold, the query was identified as “familiar” (small distance to training set) or “novel” (large distance to training set) (**Figure 2f**). The Normalized Levenshtein Distance (NLD) (see **SI Methods**) is used here as the distance metric. NLD is computed between a given query RNA and the nearest neighbor RNA (within a high-dimensional embedding space), called the explanatory sequence (**Figure 2f**). The k-nearest-neighbor (KNN) embedding graph used Euclidean distance between embedding vectors to determine neighbors (see **SI Methods**).

In scenarios where labeled training data is scarce, one option is to grow the training set iteratively by adding predicted labels of the first training round into the training set used for a second training round. The highly successful protein structure prediction model AlphaFold is one example where this strategy was employed (43). TransfoRNA uses this technique on RNAs belonging to the LOCO and NA sets and that were “familiar”, i.e., close to the training set based on their embeddings (**Figure 2g**, see below for more details).

An ensemble model was created from the predictions and explanatory sequences of individual TransfoRNA models (**Figure 2h**). The ensemble predictor collects potential explanatory sequences from the individual models for the same query. If any of the models successfully provided a relevant explanation (NLD < threshold, i.e. “familiar”), its predicted annotation would be taken as the ensemble prediction. The ensemble predictor only labeled a sequence as “novel” if none of the models could find a sufficiently plausible explanation (see **SI Methods** and **Figure S12b**).

After evaluating the TransfoRNA approach within the TCGA dataset, we trained TransfoRNA with no held-out data and used it to annotate a second dataset that was independently collected from human blood samples in a lung cancer (LC) study (8).

A more detailed description of the methods can be found in the **Supporting Online Material**. The data and TransfoRNA annotations can be found as **Supporting Online Data** and can also be interactively explored at the website www.transforna.com.

## RESULTS

### Benchmarking Transformer performance on long RNAs

TransfoRNA models use the widely spread Transformer neural network architecture for learning on irregular (non-tabular), sequential data. Other types of neural network-based machine learning, such as Convolutional Neural Networks (CNN) or Graph Convolutional Networks (GCN) could in principle also be used. TransfoRNA was therefore tested on a previously published benchmark for long non-coding RNA classification (34), where it could compete with the best CNN (nRC (44); RPX-snRC (45)) and GCN (RNAGCN (46)) alternatives (**Figure S3a**). When trained on these long RNAs (**Table S1**), however, the TransfoRNA model was not able to classify small RNAs into major classes like miRNA, rRNA, or tRNA (**Figure S3b**). It is therefore important that models are specifically trained on small-RNA data.

### Evaluating novelty prediction on held-out RNA classes and growing the training set

This small-RNA data was obtained from the public TCGA project and KBA was used to provide high-confidence (HICO) major and sub-class labels. To assess the confidence level of the model’s predictions, even when the true labels are not known, a “novelty predictor” was trained to distinguish familiar RNAs (i.e. from the same sub-classes that the models were trained on) from novel RNAs (i.e. new sub-classes that the model has not been trained on, and artificial RNAs). The term “in-distribution (ID) set” refers to data that are familiar and easy to classify (e.g. similar to training data), while an “out-of-distribution (OOD) set” contains data that the model may not easily classify, especially when coming from new and unknown (sub-) classes. Therefore, TransfoRNA first predicts whether an RNA is familiar or novel in comparison to the training data. This distinction was based on the NLD (Normalized Levenshtein Distance) novelty metric, and a threshold to optimally separate ID data from OOD data, fitted by logistic regression. TransfoRNA’s sub-class predictions were only taken into consideration when the sequences are predicted as familiar, otherwise, the predictions were discarded. While the ID set consisted of the 374 most populated sub-classes (at least eight RNA sequences per sub-class) and was further split into training and test data with a 90%/10% split, the OOD set consisted of the remaining 1549 sub-classes that had too few RNAs assigned to them in the training data, including miRNAs (one miRNA sequence per sub-class). Additionally, three sets of putative artefacts and randomly generated RNAs were included as OOD (**Figure S5a**, also see **Figure 3** and **Table 1**). This approach showed encouraging results, with high balanced accuracies (around 97% with low standard error) for sub-class models and familiar RNAs, i.e. those in the ID test set (**Figure S4a**). Except for the baseline model, AUCs of around 97% were achieved in separating the ID test set from OOD (**Figure S5b**). There was also good agreement between “familiar” non-baseline TransfoRNA annotations and low-confidence (LOCO) KBA labels (**Figure S5c**), where between 74-79% of annotations matched at least partially. For “novel” RNAs, in contrast, that match was closer to 1%. Furthermore, it was shown that major-class prediction can be fully replaced by sub-class prediction with higher performance (**Figure S4a**), likely due to the higher intra-class similarity of RNAs per sub-class compared to major class (**Figure S4b**). Therefore, only sub-class predictors were developed further. The various explored models showed differences in their annotations (**Figure S4c**) and embeddings (**Figures S4d, S6, S7**). Annotations predicted by these models (under the condition of familiarity) were used to grow the training set as outlined on the left of **Figure 3** and **Figure 2g**, leading to more training data for the final TCGA-trained models.

**Figure 3.**
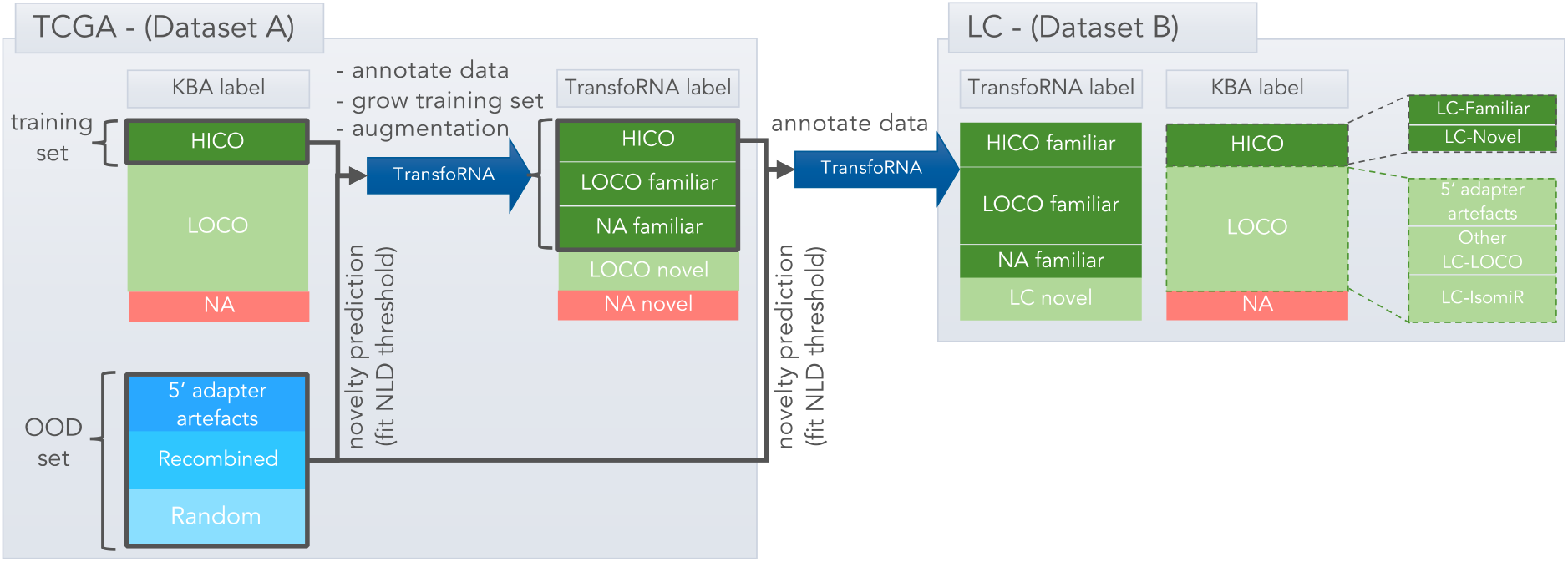
Use of novelty prediction during training and annotation. Models were trained on Dataset A (TCGA) and inference was performed on Dataset B (lung cancer study, LC). Knowledge-based annotation (KBA) was applied on both datasets. The training set consisted of all HICO annotations and those annotations considered “familiar” by the first iteration of models, all from dataset A. The LC-Familiar set (dataset B) contains all LC sequences belonging to sub-classes found in the HICO set of dataset A, making it a suitable set for gauging classification performance. A new novelty (NLD) threshold was fit to best separate the training samples from available OOD sets (artefacts, random, recombined).

**Table 1.** Different out-of-distribution (OOD) sets of RNAs, used as examples of novel RNAs to fit NLD thresholds.

| Set of novel RNAs (out-of-distribution, OOD) | category | Number of RNAs | Description |
| --- | --- | --- | --- |
| Putative 5'-adapter prefixes | real technical artefact | 1555 | The 5'-end perfectly matches the last five or more nucleotides of the 5'-adapter sequence, commonly used in small RNA sequencing. |
| Recombined RNAs | computer-generated | 200 | RNA sequences from the training set were recombined into new sequences which retain a high sequence similarity to members of the training set. |
| Random RNAs | computer-generated | 200 | Randomly chosen length and nucleotides with uniform probability. No overlap with the training set. |

### TCGA-trained TransfoRNA models can be used to confidently annotate a new dataset

To estimate transferability of a fully TCGA-trained model to new datasets, we used NGS sequencing data from a clinical lung cancer (LC) study (8). These were small RNAs observed in blood samples from patients with and without lung diseases (dataset B in **Figure 3**), as opposed to the solid tissue samples from TCGA (dataset A in **Figure 3**). The library preparation process differed partially as well, and another sequencing machine was used. Thus, this dataset contained many RNAs that were not part of the training set (**Figure 3**).

All high-confidence annotations from TCGA were used for training, without holding out any test set. Due to the ability of TransfoRNA to identify familiar RNAs in the LOCO and NA sets from TCGA (**Figure 3, S5c**), even more training samples became available, as these annotations were added as training labels (**Figure 2g, 3**). Since a part of the evaluation-phase OOD set (consisting of miRNAs and rare sub-classes of other major classes, **Figure S5a**) was added to the training set, a new novelty prediction model and NLD threshold needed to be fitted on the OOD set in **Table 1**, separately for each model. Using this learned threshold per model, annotations on the new dataset (dataset B in **Figure 3**) could be assigned a novelty (NLD) score and assigned to the “familiar” or “novel” group.

### Assessment of transferability to new datasets

To assess the performance of the TransfoRNA models, the LC dataset was divided into six distinct sets, depicted in **Figure 3**. Each of these sets served as a distinct dimension for evaluating the TransfoRNA models. KBA was used as for the TCGA dataset to assign RNAs to HICO, LOCO, and NA sets. The high-confidence (HICO) annotations within the LC dataset were separated into two categories: LC-Familiar and LC-Novel.

LC-Familiar consists of sequences belonging to sub-classes from TCGA on which the models were trained. Consequently, these sequences should ideally be predicted as “familiar” and accurately annotated with their corresponding sub-class. In contrast, LC-Novel comprises sequences from sub-classes not in TCGA, i.e. that the TransfoRNA models were not trained on. In this set, all sequences should be predicted as “novel”. The LC-LOCO set encompasses sequences categorized as low confidence based on the same predefined criteria as before (see **SI Methods**). TransfoRNA models were expected to annotate a portion of these sequences as “familiar”, thus elevating them to the HICO category (also see **Figure S5c**). The “LC-NA” set includes sequences that couldn’t be annotated by KBA. Given that HICO sequences rely on KBA annotations, it is anticipated that a smaller proportion of this set will be labeled as “familiar” by the models, in comparison to the LC-LOCO set, as shown during model development in **Figure S5c**.

The LC-IsomiRs set represents variants of the reference miRNA sequences stored in miRBase that meet specific criteria (see **SI Methods**). The LC-putative-5’-adapter set contains sequences within LC with 5’ adapter prefixes. As these sequences are considered artifacts, they should ideally be predicted as “novel.”

For comprehensive evaluation, two additional novel-by-design curated sets were included in the Total Novel set: Random and Recombined, as elaborated in **Table 1**.

### An ensemble of TransfoRNA models improved annotations

The different TransfoRNA models (except baseline) all had similar overall performance (**Figures S4 and S5**), yet still differed in their annotations (**Figure S4c**), indicating that an ensemble predictor of RNA sub-classes might perform best. Since the baseline model, as shown in **Figure S5b**, had a lower AUC in separating the ID set from the OOD sets, it was not included in the ensemble predictor. Like all other models, the ensemble model provides a sub-class and a novelty prediction for a given sequence. The sub-class prediction is considered reliable only if the sequence is predicted to be “familiar”.

The percentage of sequences mistakenly considered “novel” in the LC-Familiar set by each model is depicted in **Table 2**. The balanced accuracy of the sequences considered “familiar” is shown as well. Classification performance on the LC-Familiar set was lower for the baseline model compared to Transformer-based models. All Transformer-based models achieved almost identical balanced accuracy for sub-class prediction on the sequences considered “familiar” by each model. Although the novelty predictor also showed similar values for all Transformer-based models, except for Seq-Seq. Still, the ensemble model was able to annotate 0.9% (∼400 sequences) more when compared to the closest performer (Seq). This shows the ensemble model’s ability to annotate more sequences compared to individual models while maintaining the close performance as shown by the balanced accuracy. A confusion matrix in **Figure 4a** shows per major class that misclassifications on LC_Familiar RNAs were rare as the major class balanced accuracy was 97% (see more below).

**Figure 4.**
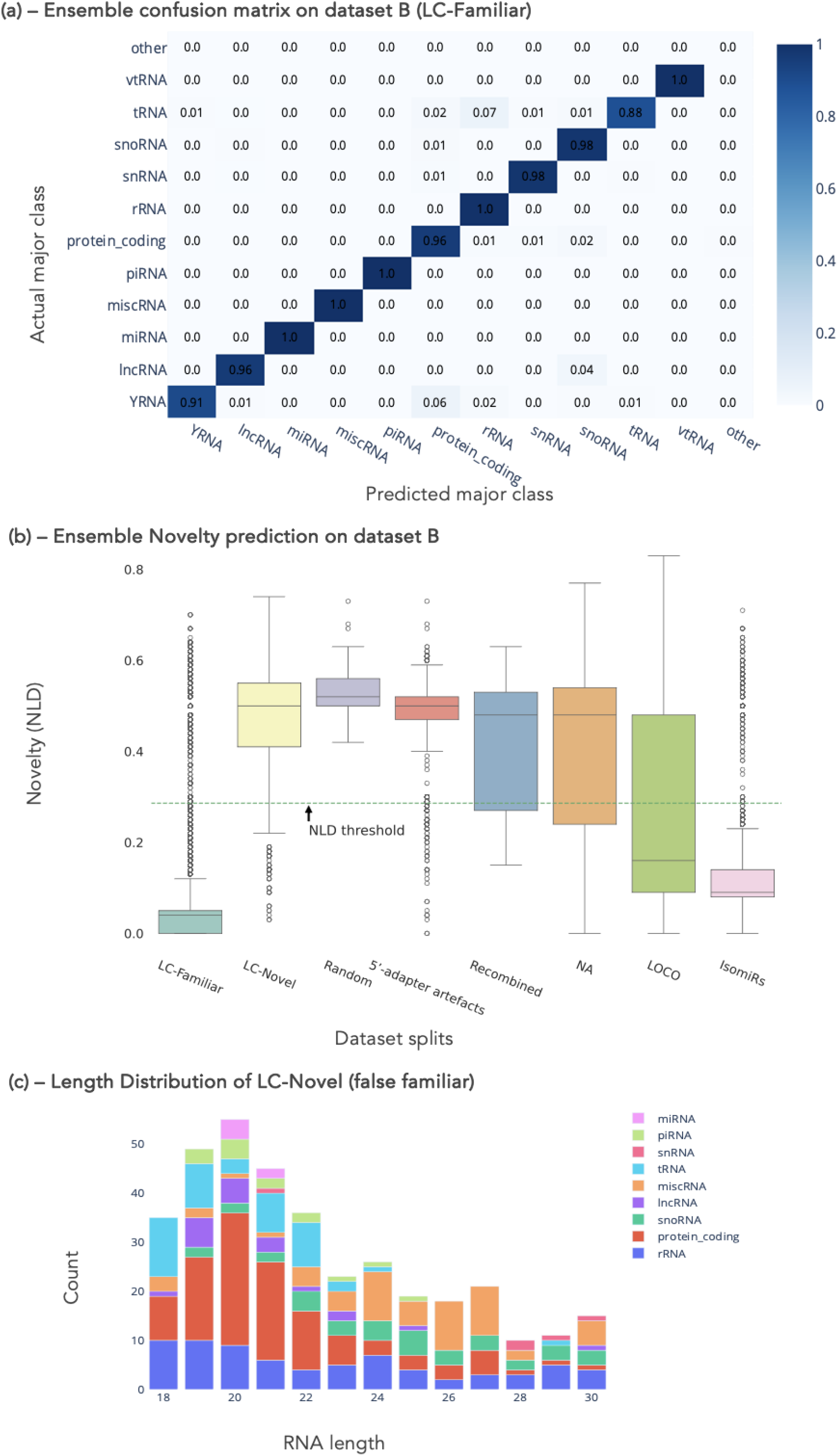
Evaluation of the fully TCGA-trained ensemble model. (a) The confusion matrix of the ensemble model was computed for the sub-class prediction of the LC Familiar set and aggregated per major class. (b) NLD Distribution for sets of sequences found in dataset B, as determined for the ensemble model. Two artificially curated novel sets were added: “Random and Recombined. LC-Familiar comprises HICO sequences found in LC belonging to sub-classes the models have seen before. LC-Novel consists of HICO sequences belonging to novel sub-classes. Putative 5’ adapters are sequences with 5’adapter prefix (could be seen as a subset of Recombined). NA and LOCO sets are sets that could not be annotated and sequences with low confidence annotations based on our internal KBA. IsomiRs are variants of the reference miRNA sequences deposited at miRBase that meet defined criteria (see SI Methods). (c) Distributions of length per major class of only those RNAs in the LC-Novel set, which were falsely predicted as familiar.

**Table 2:** LC-Familiar Evaluation. Models were trained on Dataset A (TCGA) and inference was performed on Dataset B (lung cancer study, LC). Knowledge-based annotation (KBA) was applied on both datasets. The training set consisted of all HICO annotations and those annotations considered “familiar” by the first iteration of models, all from dataset A. The LC-Familiar set (dataset B) was constructed to contain all LC sequences belonging to sub-classes found in the HICO set of dataset A, making it a suitable set for gauging classification performance. A new novelty (NLD) threshold was fit to best separate the training samples from available OOD sets (artefacts, random, recombined).

| Model | % False Novel | Sub Class B. Acc |
| --- | --- | --- |
| Baseline | 12 | 75.8 |
| Seq | 3 | 97.2 |
| Seq-Seq | 7 | 95.9 |
| Seq-Struct | 3 | 96 |
| Seq-Rev | 3 | 96.4 |
| Ensemble | 2 | 96.9 |

#### Ensemble improved novelty prediction

To evaluate the novelty prediction for the LC dataset, the NLD distribution of each of the sets was computed based on the ensemble model and plotted in **Figure 4b**. The green dashed line represents the novelty prediction threshold (NLD = 0.23). The threshold separates the model’s predictions into “familiar” and “novel” for all RNAs. Generally, the NLD distribution was as expected for each set. LC-Familiar and LC-isomiRs were mainly considered “familiar”. The LC-Novel, LC-putative-5’-adapter, Random and Recombined sets are shown to be mostly “novel” (95.0%, 97.2%, 100% and 90.8% are annotated as “novel”, respectively). As expected, the model separated the Random set further apart when compared to the other sets which have a higher overlap with the training set. For the LC-putative-5’-adapter set, the non-adapter part of a sequence may belong to a sub-class from the training set, and therefore may be close to an explanatory sequence that passes (i.e. is smaller than) the NLD threshold. Similarly, the Recombined set was made up entirely by fusing sequences from the training set and therefore had a higher chance of finding a plausible explanatory sequence.

Of the few LC-novel sequences (5% or 363 sequences) that were falsely considered as “familiar”, most were annotated as fragments of protein-coding RNAs (∼30%) or miscRNAs (∼16%) (**Figure 4c**). Protein-coding RNAs (messenger RNAs) are a large and diverse category that include a plethora of genes, splicing variants, and pseudogenes, making clear distinctions expectably difficult. A large proportion of misclassifications (40%) are made for fragments of protein-coding RNAs that map to more than 50 different genes (“hypermappers”). For misclassification of other small RNA classes, the predicted label often shares the same genomic locus as the label issued by the KBA. For tRNAs many of the false familiar labels given by TransfoRNA are isoacceptors of the KBA label, which have very similar sequences.

Because LC-NA sequences couldn’t receive annotations from KBA (i.e. the source of ground truth for TransfoRNA), it is anticipated that a smaller proportion of them would be categorized as “familiar” compared to LC-LOCO, where KBA annotations were feasible but sequences were labeled as low-confidence due to other factors (see **Figure 2c, SI Methods**). The ensemble annotated 30% of the NA set that could not be annotated using the KBA.

#### Accurate identification of isomiRs

So far, a very strict definition of miRNAs had been used, where each miRNA type was represented by a single sequence which is listed as the canonical “reference” sequence at miRBase (refmiR). The TCGA-trained TransfoRNA models have thus only encountered a single example sequence per miRNA (each a different sub-class), and only 594 in total. Within the LC dataset, 6849 isomiRs (isoforms of each refmiR) were largely identified correctly (**Table 3**). The ensemble model marks fewer of the isomiRs as “novel” (3%), compared to the individual models (12% and higher). The balanced accuracy of the ensemble model on the prediction of isomiRs is 98.8% (sub-class prediction).

**Table 3.**
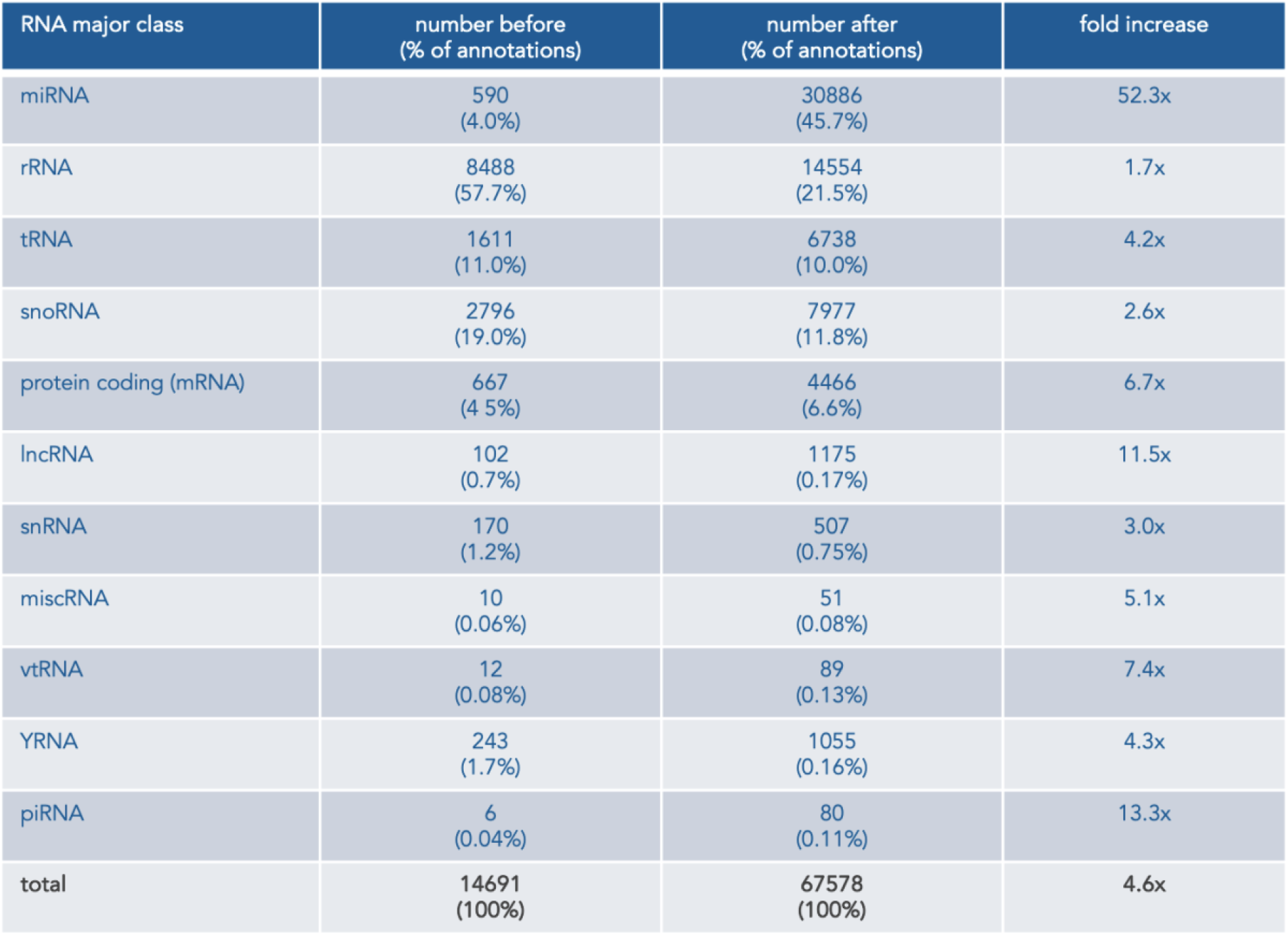
Number of high-confidence annotations before and after using TransfoRNA ensemble model (TCGA)

#### Extended high-confidence annotations within a new dataset

The performance of each individual model in annotating sequences with high confidence was illustrated in **Table 3**, alongside the ensemble model. Across all sets, the ensemble model outperformed each individual model. This suggests that each individual model had developed its own subfield of expertise which was bolstered through an ensemble. Further details are available in **Figure S9**, which lists the additional number of sequences annotated by the ensemble model within each major class. In the case of all LC sequences (comprising ∼208k sequences), not including the LC-Novel set, the ensemble model demonstrated a 5% increase (18.7k sequences) in annotations compared to the closest Transformer-based single models (Seq-Seq, Seq-Struct and Seq-Rev). When contrasted with KBA annotations, TransfoRNA achieved confident (“familiar”) annotations for 62% (∼82k sequences) of the LOCO set and 30% (4.7K sequences) of the NA set in dataset B. However, with regards to LC-Novel, a drawback of consolidating knowledge from individual models is the aggregation of false positives. The ensemble model mistakenly designated 143 more sequences as HICO in comparison to the best-performing single model. However, this may still be considered a negligible number. When mapping the sub-class predictions of the LC-Novel set to their respective major class only a low balanced accuracy was achieved, confirming that the novelty prediction is essential as a pre-filter for annotations (**Figure S11**).

#### More high-confidence annotations added to the TCGA dataset

The percentage of HICO annotations in dataset A (TCGA) could be increased from 19.6% to 69.6% by adding TransfoRNA predicted annotations to the original KBAs (**Figure 5**). In the right pie chart, sequences were assigned a LOCO or HICO label for a given model according to their assignment as novel or familiar, respectively (**Figure S12a**). The largest-fold increase (**Table 4**) was observed for miRNAs (52.3 fold) where TransfoRNA was able to classify ∼99% of isomiRs into the right sub-class, based on just a single example for each. For most other major classes, HICO-annotated sequences increased by more than two-fold. rRNAs were already highly covered by the KBA and augmentation approach and only increased slightly. The progression of the number of HICO sequences in TCGA during the evaluation and testing phase is available in **Figure S10**. All TransfoRNA annotations used herein are part of the **Supplementary Online Data** and www.transforna.com.

**Figure 5.**
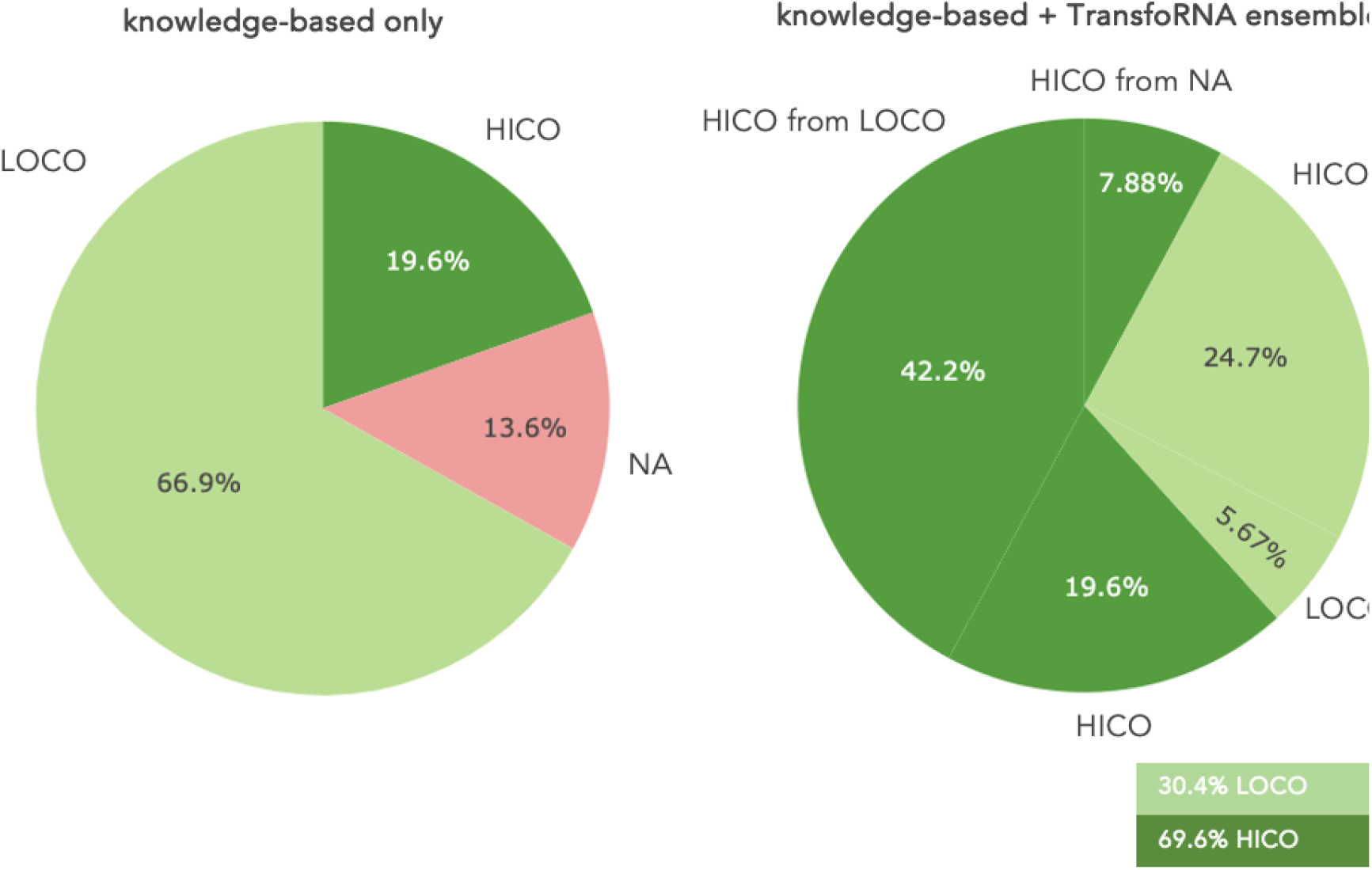
Quantification of high-confidence annotations before and after applying the production version of TransfoRNA within the full TCGA dataset. (a) Comparison of knowledge-based annotation (KBA) and TransfoRNA-ensemble annotations regarding confidence levels, annotation coverage, and sources of new high-confidence (HICO) and low-confidence (LOCO) annotations after applying TransfoRNA to high-confidence, knowledge-based training data from TCGA. The HICO set increased 50.0% while the LOCO set decreased by 36.5% of the whole data. The original NA set (no annotation) ends up in either

### Visualization of all RNAs with high-confidence annotations

Each model that is part of the ensemble encodes RNA sequences in different ways as embeddings. **Figure 6** shows these 100-dimensional (or 200-dimensional where two inputs are combined) embedding spaces via UMAP dimensionality reduction (47) as 2D scatter plots. The three sequence-only TransfoRNA models (Seq, Seq-Seq, Seq-Rev) are most notable for their dense clustering of similar isomiRs (miRNAs) and strand-like clusters for consecutive rRNA sequence bins. Endowing the models with structural information (Seq-Struct) also changes the priority for clustering, grouping RNAs more by their structural properties, such as the degree of base-pairing.

**Figure 6.**
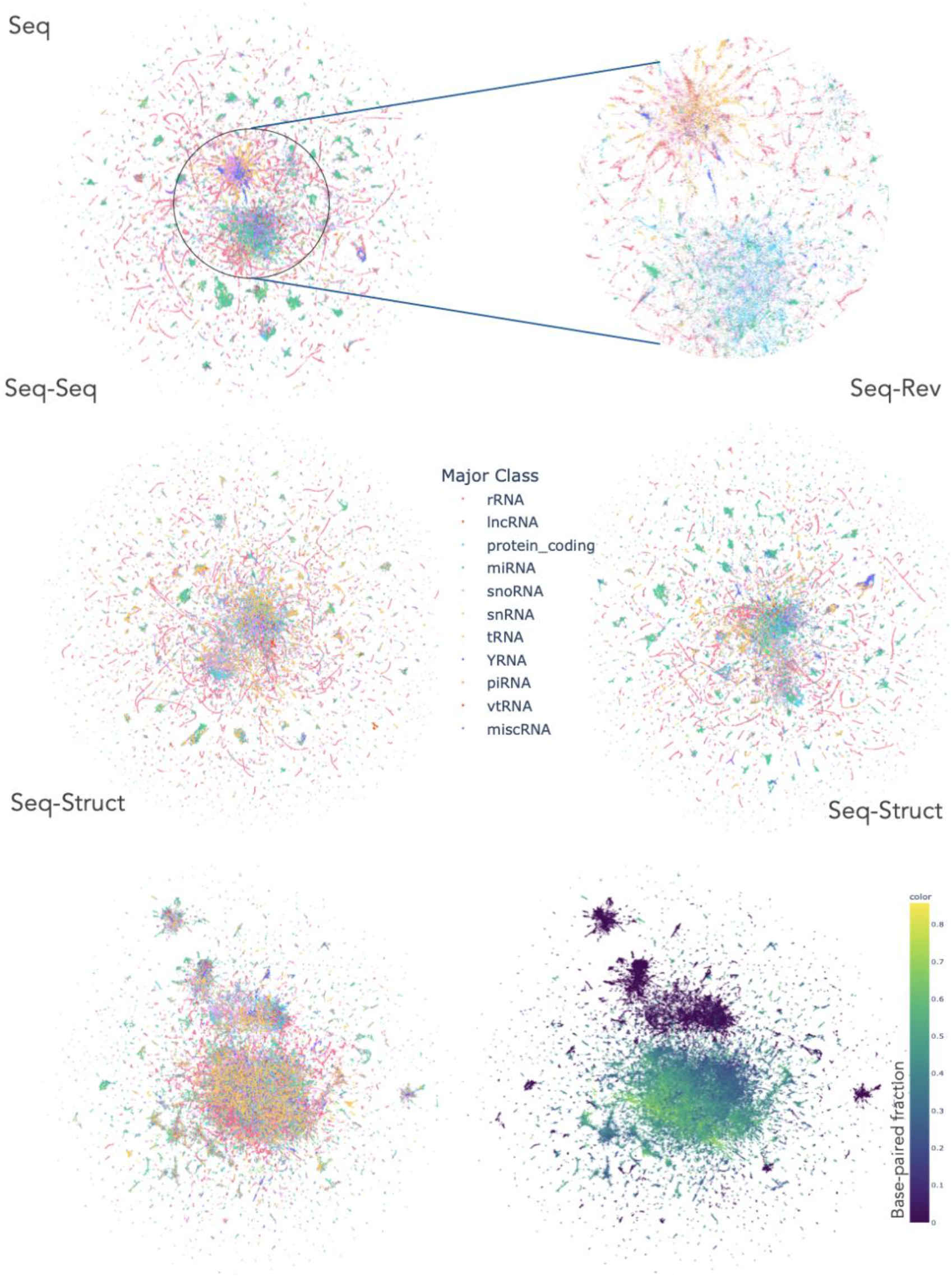
RNA embedding spaces created by different TransfoRNA transformer based models. Each dot in these UMAP plots represents an RNA converted into embedding vectors and subsequentially mapped to a 2D space. Distances in the UMAP imperfectly reflect distances in the high-dimensional embedding space. Colors correspond to major class labels as indicated. For the Seq model, the cluttered region at the center is magnified. The base-paired fraction is the ratio of base-paired nucleotides over the length of the sequence. All RNAs with high-confidence annotations, from KBA or predicted by TransfoRNA, from either dataset (TCGA, LC) are shown.

## DISCUSSION

We have demonstrated an innovative annotation workflow for small RNA sequencing data that combines high-confidence ground truth obtained through conventional mapping methods for a small number of RNA sequences with AI-assisted extrapolation to any other RNA with a built-in measure of annotation confidence. TransfoRNA models work very well to preserve the small set of already reliable knowledge-based annotations, as indicated by a high classification accuracy on an in-distribution test set (**Figure S4a**) and new RNAs in another dataset (**Figure 4a**). However, it takes a different approach when it comes to annotating most other small RNA sequences, namely those that do not precisely match the reference sequences and where multiple alternative annotations are possible. Enforcing a general set of annotation rules for precedence and mismatches is likely to work well only in some cases, but not in others. Instead, TransfoRNA extrapolates automatically from a small set of annotations, based on the vectorized representations of RNA sequences (embeddings) that it has learned.

A key contribution was to design a metric of classification certainty of TransfoRNA models that measured how close a new RNA was to those that the model had already seen annotations for: the normalized Levenshtein distance (NLD). A learned threshold for this metric served as a novelty predictor to identify RNAs below the threshold, for which the model can make a confident annotation. The remaining (novel) RNAs above the threshold may be excluded from downstream analysis or analyzed separately. For example, RNA expression levels, where available, could be used to remove potential technical artefacts which are more likely to have low expression and occur in fewer samples.

TransfoRNA also demonstrated excellent generalization ability by providing high-quality annotations on a second, fully independent dataset, which contained a large number of new RNA sequences. Annotations were close to perfect for new members of the already trained-on RNA sub-classes.

Given the high interest in miRNAs, we have taken a very conservative approach here by initially only considering the miRBase reference sequences as high-confidence annotations. Therefore, when training TransfoRNA on the entire TCGA set (**Figure 3**), only 594 annotated miRNA sequences were used as ground truth labels. Still, more than 30000 isomiRs could be recovered among the remaining RNAs in TCGA, covering 99% of putative isomiRs. Models were not re-trained on predicted miRNA annotations, as was done for other classes. Future versions of TransfoRNA could therefore explore multiple iterations of growing the training data from the most confident predictions (**Figure 2g**). The 75080 small RNAs obtained from TCGA are still a relatively small dataset, not covering all known miRNAs. TransfoRNA models have therefore not reached their full potential scale yet, which could be achieved by including more NGS data.

A major benefit of using actual RNA reads from NGS as training data, instead of just genomic reference sequences, lies in the inclusion of real sequence variants as they occur in the biological samples analyzed. The iterative approach of growing the training set from confident predictions ensures that these variants are also incorporated into the models. As mentioned above, more growth iterations could be done in principle. The larger the training corpus, the more genetic variation present in human populations will be covered. Applying expression filters on the training data is important to exclude most (rare) technical variants due to incorrect base-calling.

Several notable findings were made around the machine learning setup described here. First, a more detailed sub-categorization of the major RNA classes facilitated the classification task considerably. These sub-classes carry additional biological information that would otherwise have to be learned by the model instead. For example, there is little commonality among rRNA fragments so that the model must learn the explicit mapping of each fragment to the “rRNA” major class. An rRNA sub-class on the other hand is defined in such a way that similar sequences are grouped in narrow bins (i.e. as a common region along one of the full-length rRNA molecules).

Second, all Transformer-based models performed significantly better than the simple baseline model. Transformers have the ability to learn patterns and connections between any positions along the full RNA sequence that simpler models don’t. Future work should focus on explainability methods to highlight patterns in the inputs responsible for a particular classification outcome. The representation of the inputs clearly mattered and the different TransfoRNA variants made partially different predictions, but it is not clear yet, how these differences came about in detail. The ensemble-model approach could successfully combine these different model variants and enhance performance further.

Future work should also extend the augmentation approach used here to deal with the limited expression of fragments derived from larger RNAs. These can make up a large part of the data (e.g. rRNA) but not all parts of parental RNA are found or covered equally. A particular challenge might come from protein coding RNAs and alternative splicing variants which might contribute to a large portion of rare small RNAs but might be difficult to annotate. Current shortcomings in annotating major classes among novel RNAs (see above) can be addressed in the future.

Expression data may also be included to resolve annotation uncertainty as well as information on the sample type or tissue of origin. Other types of omics data may also be incorporated. This is important, because due to their short length, some small RNAs will remain ambiguous if they are identical to precursors from different classes.

As a machine learning model, TransfoRNA can be easily trained on other sequencing datasets (e.g. with longer read lengths) and extended as new ground truth becomes available. Future work should also investigate the potential for transfer learning, where a pretrained model, e.g. from one organism, is fine-tuned on little additional data to make annotations for a different organism.

An additional advantage of TransfoRNA are the embeddings that map each sequence to a continuous space which can be visualized in a UMAP plot (**Figures 6, S4d, S6, S7**). These embeddings contain potentially rich information that may be used for various other data analyses. Each model provides different embeddings with different properties. Seq-Struct, for instance, groups RNAs more by their structural properties and so on. Embedding plots can also be a useful tool for exploring the neighborhoods of RNAs, identifying clusters of similar RNAs, and finding similarities between RNAs from different classes.

As a conclusion, it is important to note that accurate and exhaustive annotation of small RNAs remains a challenging task with room for further improvement. Knowledge-based annotation remains an important prerequisite, and more research must be invested in checking and expanding the validated training labels used for AI-based techniques. Nevertheless, we believe that TransfoRNA makes several important contributions towards AI-based small-RNA annotation.

## ACKNOWLEDGEMENTS

The results published here are in part based upon data generated by The Cancer Genome Atlas (phs000178.v11.p8.c1) managed by the NCI and NHGRI. Information about TCGA can be found at http://cancergenome.nih.gov. We thank Dr Phoebe Chan for access to long non-coding RNA benchmark data.

