## Supplementary Information for "TransfoRNA: Navigating the Uncertainties of Small RNA Annotation with an Adaptive Machine Learning Strategy"

### SI METHODS

#### Hardware and Software

The proposed architectures were implemented using the PyTorch (https://pytorch.org) and Skorch (https://skorch.readthedocs.io/en/stable/#) Python libraries. The open-source implementation of ConveRT's architecture (https://github.com/jordiclive/Convert-PolyAI-Torch) was used as a codebase for building the models. The training was performed using Nvidia RTX A6000 GPUs (each with 48 GB of memory) and an AMD Ryzen Threadripper 3990X 64-Core Processor.

#### RNA-seq small-RNA datasets

The Cancer Genome Atlas (TCGA) (6) is a landmark cancer genomics dataset comprising sequenced and molecularly characterized expression data of over 20000 primary cancers and matched non-diseased tissue samples from 33 cancer types. The small RNA sequencing data was described earlier (7).

A second dataset was independently obtained by sequencing clinical blood samples from lung clinic patients as described previously (8). Different protocols were used for sample collection and library preparation.

### Data preprocessing

Raw sequencing data of 11041 miRNA-Seq measurements was downloaded from the GDC Data Portal (https://portal.gdc.cancer.gov/) with permission. A read count matrix was generated for the 75080 sequences that have a minimum length of 18 nucleotides and that are present in at least 30 samples of each associated cancer/tissue type.

### Small RNA annotation using published tools

To annotate the 75080 small RNA sequences, we used the rRNA- and tRNA-focused small RNA annotation tool SPORTS (9) (version 1.1; parameters: -M 2) and the more miRNA-focused annotation tool UNITAS (10) (version 1.7.7; parameters: -species_miR_only) together with their pre-compiled annotation databases and the snoDB resource v2.0 as additional snoRNA reference (11). Based on the annotation matrices of the two approaches each sequence was assigned to the following major classes: miRNA, rRNA, tRNA, lncRNA, snoRNA, snRNA, miscRNA, vtRNA, YRNA, piRNA, protein_coding (see **Figure S1a**). Sequences that could not be assigned to these classes are referred to as the "no annotation" (NA) set.

### Small RNA annotation with increased annotation coverage

As both annotation tools map the sequences sequentially to the different small RNA class databases, the order of mapping determines the annotated small RNA class. However, small RNA sequences can have not only one but several potential precursors. In sequential annotation pipelines these other potential parental RNAs belonging to small RNA classes of a lower mapping hierarchy are neglected, even though a mapping hit could have less mismatches. To overcome this, we generated one extensive database of human transcripts that contains the miRNA hairpins from miRBase ((12); depleted for duplicates at other genomic loci), the mature tRNA sequences from GtRNAdb (13) extended for leader and trailer sequences by adding the 50 nucleotides upstream and downstream of the mature tRNAs), the rRNA and YRNA sequences from NCBI (Sayers et al. 2022), the snoRNA and scaRNA sequences from snoDB (Bouchard-Bourelle et al. 2022), the snoRNA and scaRNA sequences from snoDB (11), the piRNA cluster sequences from piRNA cluster DB (15), as well as the lncRNAs, snRNAs, vtRNAs, sRNAs, MT-tRNAs, MT-rRNAs, misc_RNAs and cDNA sequences from ENSEMBL (16) (depleted for lncRNAs that are miRNA host genes). Using bowtie (17) (parameters: -a --norc), we first mapped the small RNA sequences against this reference database, prohibiting any mismatches. Sequences without perfect match were next mapped to the database again allowing for one mismatch this time. Sequences that did not map once more, were indented to map allowing for two mismatches and in unsuccessful cases a last time allowing for three mismatches.

Small RNA sub-classes were defined in the following way. For miRNAs, the canonical name of the reference sequence was used for all associated isomiRs that follow a set of certain rules that are based on the latest isomiR research (18). Hence if the sequences map to the precursor hairpin with an overlap to a mature miRNA in the range of +/- 2nt. Apart from non-templated 3′ poly-A or poly-T additions, only A>G or C>T in seed region were allowed mismatches. In cases where these rules are violated, or for mappings to other parts of the hairpin the name of the hairpin was used (see **Figure S1b**). For tRNA-derived small RNAs, the sub-class names are a composite of the three-letter amino acid code and the anticodon of the full tRNA. tRNA-derived small RNAs that match the 5′ end of the mature tRNA are categorized as 5p-tR-halves if their length is bigger than 30 nucleotides otherwise they are considered 5p-tRNA fragments (same rules apply for RNAs matching the exact 3′ end of the mature tRNA). For all other small RNA classes, sub-classes were defined for ~25-nucleotide bins along the sequence for each parental RNA separately. Sequences with more than 50 potential mapping references were labeled as hypermappers.

**Augmentation of under-represented RNA sub-classes**

Certain small RNA sub-classes are represented with too few sequence variants in the training dataset to allow profound learning. To enhance coverage, an augmentation strategy is employed wherein size-appropriate segments from the precursor of the underrepresented sub-class were added to the training set. Within this approach, sequences are drawn from a specific bin of the precursors, spanning from position 'x' to position 'x + bin length'. To maintain alignment with the training set's sequence length distribution, sequences of 18 to 30 nucleotides are sampled from the bin. Since the binning positions are arbitrary, RNA fragments sequenced through NGS could lie within a bin but could also overlap with neighboring bins. To account for this possibility, sampling incorporates the neighboring bins, however, ensures that the annotation of a sequence matches the bin with which it has the highest overlap.

### Putative artefacts with 5′-adapter prefixes

Small RNA sequencing requires to append sequencing adapters to both ends of the biological sequence of interest. As the sequencing process typically starts right after the 5′ adapter, the reads do not contain the 5′-adapter sequences, but start with the sequence of interest right from the start. However, we identified a small number of sequences that appear to have remnants of the 5′ adapter at their 5′ end. This adapter, or variants thereof, was used in the TCGA libraries (5′-GTTCAGAGTTCTACAGTCCGACGATCTGGTCAA-3′) as well is in the previously published LC blood sample dataset (8).

### Recombined training set sequences

Each recombined sequence was constructed from two sequences found in the training set and belonged to two distinct sub-classes. The first sequence was truncated between 60 and 70 % of the sequence and then attached to 30 to 40 % of the second sequence.

### Random sequences

Random sequences were generated, one nucleotide at a time, from a uniform distribution of four possible outcomes; A,C,T and G. The length of each sequence is sampled randomly from 18 to 30 nucleotides, corresponding to the minimum and maximum sequence lengths in the training set.

#### Definition of high confidence (HICO) annotation

While the goal of the initial annotation was to annotate as many small-RNA sequences as possible, we defined a subset of high-confidence annotations (see **Figure S1d**) for training of our TransfoRNA models. We defined sequences as high-confidence if they mapped to their precursor transcript without mismatches, insertions, or deletions. The set of high-confidence annotations was further depleted for sequences with putative 5′ adapter prefixes and sequences that mapped to bacterial or viral RNA genomes (NCBI Refseq data) or genomic sites of transposable elements. Furthermore, only sequences with a single major and sub-class annotation were considered. High-confidence miRNAs additionally were required to perfectly match the mature reference sequences deposited in the miRBase database of release 22.1 (12) (see **Figure S1c**).

#### Sequence Alignment similarity

Pairwise sequence similarities shown in **Figure S4b** is computed based on the "pairwise" Biopython package (https://biopython.org). Given any two sequences, the score is computed while setting a score of zero for mismatches and without penalizing end gaps or extended gaps. "Within class" refers to computing the scores of all sequence pairs sharing the same class while for "between class" sequences belonged to different classes. This was done for both major and sub-class and each score is normalized by the length of the longest sequence amongst the two sequences, mapping the scores in the range of 0 to 1.

**General TransfoRNA model**

*Sequence Representation.*

A sequence $j$ with length $l$, $S_j^l, \{s_1, s_2, s_3 .. s_l\}$, is tokenized by sliding a window of size $\boldsymbol{w}$ one nucleotide per step resulting in a sequence of tokens, $T_j^{l-w}$ of length $l - w$. For instance, the sequence ACTTGA is converted to three tokens $(t_1, t_2, t_3)$; ACTT, CTTG and TTGA. Each token is assigned a unique integer identifier. All unique tokens $\boldsymbol{U}$ in the dataset are obtained by tokenizing all sequences in the training set. **Figure S2-a-I** shows an example of a sequence tokenized with window size $w = 1$.

*Noise.*

Following Godwin et al. (19), a regularization module ("noisy tokens") adds noise to the input tokens for generating more robust embeddings. Unlike Transformers (20), which work on a continuous feature space, the noise module operates on a discrete space of token ids. Tokens of sequence j with length k, $T_j^k$ are flipped randomly with a 10% probability. The probability distribution of a random token, $T^i$, at position $i$ to assume its original value, $t^i$ against assuming a different value, $d$ is:

$$P(T^i = t^i) = 0.9$$

$$p(T^i = \{d : d \in U, d \neq t^i\}) = 0.1$$

*Token Encoder.*

Given the noisy token ids, the token encoder converts ids to a continuous vector space of dimension $D_{in}$ following (20). In parallel, the positional encoder introduced by ConveRT (21) is implemented to compute a unique continuous vector space of the dimension $D_{in}$ for every unique position in the sequence, independent of the values of the input tokens. Finally, the token embeddings are added to the positional embeddings to form the position-aware embeddings, $\boldsymbol{X}$. Adding the positional embeddings to the token embeddings allows the model to discern tokens at different input positions.

*Self-attention.*

The position-aware embeddings $X \in R^{T*D_{in}}$, where T is the number of tokens, are passed to the self-attention module from Transformers (20), where the attention scores $A_s$ between every combination of tokens is computed. $\boldsymbol{A_s}$ is obtained by computing the dot product between the queries $Q$ and keys $K$. Where $K = W_k * X$ and $Q = W_q * X$. Both $W_q$ and $W_k$ are learnable weight matrices represented by a linear feed-forward layer, depicted in **Figure S2-a-II**.

*Equation 1*

$$\mathrm{A_s} = X\mathrm{W_k}\mathrm{W_q}^T X^T$$

$A_s$ has the dimension of $T * T$, where a given entry of $A_s$ at position $i, j$, $a_s^{i,j}$ reflects the attention score between that token at position $i$ ascribes to the token at position $j$. Since $i$ and $j$ are bound from 1 to $T$, this ensures that there is an attention score between every token combination in both ways, thus, ensuring minimal prior knowledge incorporation.

A Softmax layer is then applied on top of the attention scores scaled by the square root of the token embedding dimension, $D_{in}$. The scaling factor prevents large values from dominating which could prevent reasonable gradient magnitudes from backpropagating through the network (20). The output of the SoftMax can be viewed as attention probabilities which are used to update the embeddings of each token. The new token representation $X_{new}$ is then obtained by multiplying the attention probabilities by the value matrix $V$, where $V = X * W_v$. This can be depicted in the following equation:

*Equation 2*

$$SelfAttention = X_{new} = \mathrm{Softmax}\left(\frac{\mathrm{A_s}}{\sqrt{D_m}}\right) * \mathrm{V}$$

The learnable matrices $W_q, W_v$ and $W_k$ have equal dimensions of $D_{in} * D_{out}$. Throughout all tasks, $D_{in}$ is set to be 512 whereas $D_{out}$ is set to be 64. The number of tokens $T$, however, is task-dependent and is set to the maximum sequence length in the dataset. The value of T is depicted along with every task in the Results section. It is worth noting that the keys, queries, and values $\{K, Q, V\}$ are fully-connected layers that are applied to the embedding of each token in parallel which makes training efficient.

To improve the quality of the gradients passed through the network, residual connections are added (22). Concretely, both the input of the self-attention module ($X$) as well as its output ($X_{new}$) are first added and then passed to a normalized layer (22). To allow this, the normalization and self-attention layers have the same output dimensions, $D_{out}$.

*Multi-headed attention.*

Single-headed attention is a special case of the multi-headed attention (MHA). Each head $i$ receives the same triplet matrices $\{Q, K \; and \; V\}$ which are then multiplied by $\{W_i^q, W_i^k, W_i^v\}$ respectively as shown in **Figure S2a-III**. Each head then computes the self-attention operation in parallel where the outputs of all heads are then concatenated to produce the final Attention score.

*Equation 3*

$$head_i = SelfAttention(QW_i^q, KW_i^k, VW_i^v)$$

*Equation 4*

$$MHA = concat_{h \in nheads} \, [head_1, head_2, .., head_h] \, W_{out} + b_{out}$$

Where $W_i^q, W_i^k, W_i^v \in R^{D_{in} * D_{out}}$, $W_{out} \in R^{hD_{out} * D_{in}}$ and the bias, $b_{out} \in R^{hD_{out} * 1}$. While each of the multi-headed attention heads preserves almost the same information (23), it still helps to encode multiple relationships and nuances for each input token.

*Reduction.*

Following the Universal Sentence Encoder (24), we convert the learned representations to a reduced fixed length embedding vector, $\mathrm{R}^{\mathrm{s}}$, by adding the embedding representations of all tokens of a given sequence $S$, then scaling it by the sequence length: $\mathrm{R}^{\mathrm{s}} = \left(1/\sqrt{\mathrm{T}}\right) * \sum_{\mathrm{t}=1}^{\mathrm{t}=\mathrm{T}} E_{\mathrm{t}}^{\mathrm{s}}$, where $E_t^s$, is the embedding of token $t$ belonging to sequence $S$.

*Specific models with different inputs.*

RNA sequences have other related properties such as secondary structure and expression. Incorporating such multimodal input in the training requires designing architectures that are suitable. Moreover, sequences can be represented in various ways which might affect the performance of the models. To this end, we chose to design models that can learn from various sequence representations as well as different input modalities. All six models introduced are depicted in **Figure 2d and S2b**.

*Baseline model.*

To gauge the utility of TransfoRNA, a simple baseline is proposed. The sequence tokens are fed to an embedding layer that converts each token id into a continuous vector representation. All token embeddings are then concatenated and represented as $E^{\mathrm{seq}}$ in **Figure S2b**. $E^{\mathrm{seq}}$ is then fed to a linear layer that outputs the predictions over all classes. Cross entropy loss is then used to backpropagate the error between the true and predicted classes. The output of the classification layer is either set to the number of sub-classes or major classes depending on the task.

*Seq TransfoRNA model*

To gauge the effectiveness of designing various sequence representations, a straightforward representation is employed as a baseline. Sequences are tokenized and fed to the TransfoRNA block as shown in **Figure S2b**, where the reduced representation $R^{\mathrm{seq}}$, is used as an input to a linear layer.

*Seq-Seq TransfoRNA model.*

Sequences are tokenized and split into two sets of tokens; one set with the sequence tokenized starting position zero and the other skips the 0th position and tokenizes the sequences, starting at position 1.

As shown in **Figure S2b**, Seq-Seq utilizes two TransfoRNA blocks to learn from the two reduced representations of sequences. The tokenization is done with an overlapping window, therefore both TransfoRNA blocks in Seq-Seq take almost the same sequence (apart from the first and last token that might be present/missing in one or the other sequences). However, each sequence would be tokenized differently, hence, the models aim becomes learning a sequence representation capable of assigning similar embeddings to sequences tokenized differently due to a shift in the start position.

*Seq-Struct TransfoRNA model.*

Secondary structure is a simplified approximation of the actual 3D structure, only focusing on pairwise molecular binding (attraction) between nucleotides, often between sequence positions far apart. One common representation of secondary structure is the "dot-bracket" notation where each position along the sequence is mapped to one of a few characters, most frequently "(" and ")" for matching base-pairs and "." for no pairing. See the Vienna RNA software package (25) for examples. Note however that these representations are predictions and not directly based on experiments.

To test whether RNA secondary structure contains valuable information correlating with the RNA class, a model that learns form multimodal input is implemented. The sequence $S^{seq}$ and its corresponding secondary structure $S^{struct}$ are passed into a separate TransfoRNA as shown in **Figure S2b**. The sequence's reduced embedding representation is computed as $R^{\text{seq}}$ and $R^{\text{struct}}$ are concatenated and fed to a linear layer.

*Seq-Rev TransfoRNA model.*

The positional encoder creates a unique embedding vector per each token position which is incremented along with the token position inducing an inductive bias that sequences are read left to right. Seq-Rev adds an additional bias where sequences can be also read in the opposite direction by reversing the sequence and applying the same positional encoder. The sequence and its reverse are then fed, each into a separate TransfoRNA block that output $R^{seq}$ and $R^{rev}$. Both reduced embeddings are then concatenated and fed to a linear layer.

*Ensemble TransfoRNA model.*

Predictions from all TransfoRNA models were used in the construction of the ensemble predictions. There are three cases: a) If only one of the models can predict a query sequence as "familiar", meaning that the model can capture a relevant explanatory sequence, then its prediction of that model is adopted by the ensemble model; b) If more than one model can predict a query sequence as "familiar", then the prediction of the model with the least NLD is considered; c) If none of the models could predict a sequence as "familiar", then the ensemble predicts the query sequence as "novel". A graphical illustration of the possible permutations is available in **Figure S12b**.

*Graph Creation*

To create a k nearest neighbor graph from the sequence embeddings, the nearest neighbor method offered by Scikit-learn (26) was used. The algorithm employed for the graph creation was the brute-force search. Minkowski distance with l2 distance was used as a distance metric which resolves to the Euclidean distance.

*Training settings.*

Cosine annealing (27) is employed as a learning rate scheduler. This scheduler starts with a large learning rate which is then rapidly decreased to a minimum value before being increased rapidly again, forming cycles of rapid increase followed by rapid decrease. The maximum and minimum values are fixed throughout all tasks to $\eta_{min} = 1e^{-3}$ and $\eta_{max} = 1$.

The training sets are stratified and split randomly into 90% and 10% to obtain training and test sets. 10% of the training set is then used for validation. Sequences are tokenized with a window size of 2 and percentage of noisy tokens are set to 10%. Following that, training is done with a maximum of 3000 epochs which is chosen based on observing minimal improvement on the validation set. Model parameters at the epoch performing the best on the validation set is saved and used for testing. This process is repeated five times for each model.

*Hyper-parameter tuning.*

The Initial hyper-parameter settings were optimized based on the external benchmark presented in **Table S2**.

*Novelty prediction training.*

A univariate logistic regression is used for training ID vs OOD where both are combined and randomly split in a stratified manner into training, validation, and test sets with respective proportions of 80%, 10% and 10%. OOD vs AA classifier is trained in a similar manner, however, the model used is a linear SVM.

**Losses and metrics**

*Cross Entropy Loss.*

Models in the TransfoRNA framework are trained in a supervised manner for predicting separately RNA major and sub-classes. The loss described as the difference between the logits (class predictions) and the true labels is computed based on a multi-class cross-entropy loss which is employed to update the parameters of the whole model.

Multi-class cross entropy loss:

*Equation 5*

$$\mathcal{L}_{CE} = -\sum_{i=1}^{n\ classes} y_i \log(\hat{y}_i)$$

Where $y_i$ is a one-hot encoded class label and $\hat{y}_i$ are the predicted probabilities for all classes.

The logits reflecting the model's confidence that a given RNA belongs to each of the classes is used as input for the novelty prediction.

*Area under the ROC curve.*

AUC-ROC is a metric used to measure performance of a model under various threshold settings and is computed using the TPR (true positive rate) and FPR (false positive rate):

$$TPR = \frac{TP}{TP + FN}$$

$$FPR = \frac{FP}{FP + TN}$$

True negatives $TN$, true positives $TP$, false negatives $FN$ and false positives $FP$ are computed using the confusion matrix.

*Accuracy.*

Accuracy provides a measure of how close model predictions are to the true labels and is computed as:

$$ACC = \frac{TP + TN}{TP + FP + TN + FN}$$

*Balanced Accuracy.*

The balanced accuracy (bACC) is used as a performance metric when classes are unbalanced. It is computed using the true positive rate (TPR) and the true negative rate (TNR) as follows:

$$bACC = \frac{TPR + TNR}{2}$$

*Normalized Levenshtein distance (NLD).*

Levenstein distance is a metric used to measure the proximity between two strings. It is thought of as the minimum number of changes, that takes the form of deletions, insertions or substitutions, to make one sequence identical to another. It is formulated as:

$$lev(s_a, s_b) = \begin{cases} |s_a| & if\ |s_b| = 0, deletion \\ |s_b| & if\ |s_a| = 0, insertion \\ lev\big(tail(s_a), tail(s_b)\big) & if\ s_a[0] = s_b[0] \\ 1 + min \begin{cases} lev(tail(s_a), s_b) \\ lev\big(s_a, tail(s_b)\big) \\ lev(tail(s_a), tail(s_b)) \end{cases} & otherwise \end{cases}$$

To get the Normalized Levenshtein Distance, NLD, the Levenshtein distance is divided by the longest sequence as follows:

$$NLD(s_a, s_b) = \frac{lev(s_a, s_b)}{\max(\{|s_a|, |s_b|\})}, \quad [0,1]$$

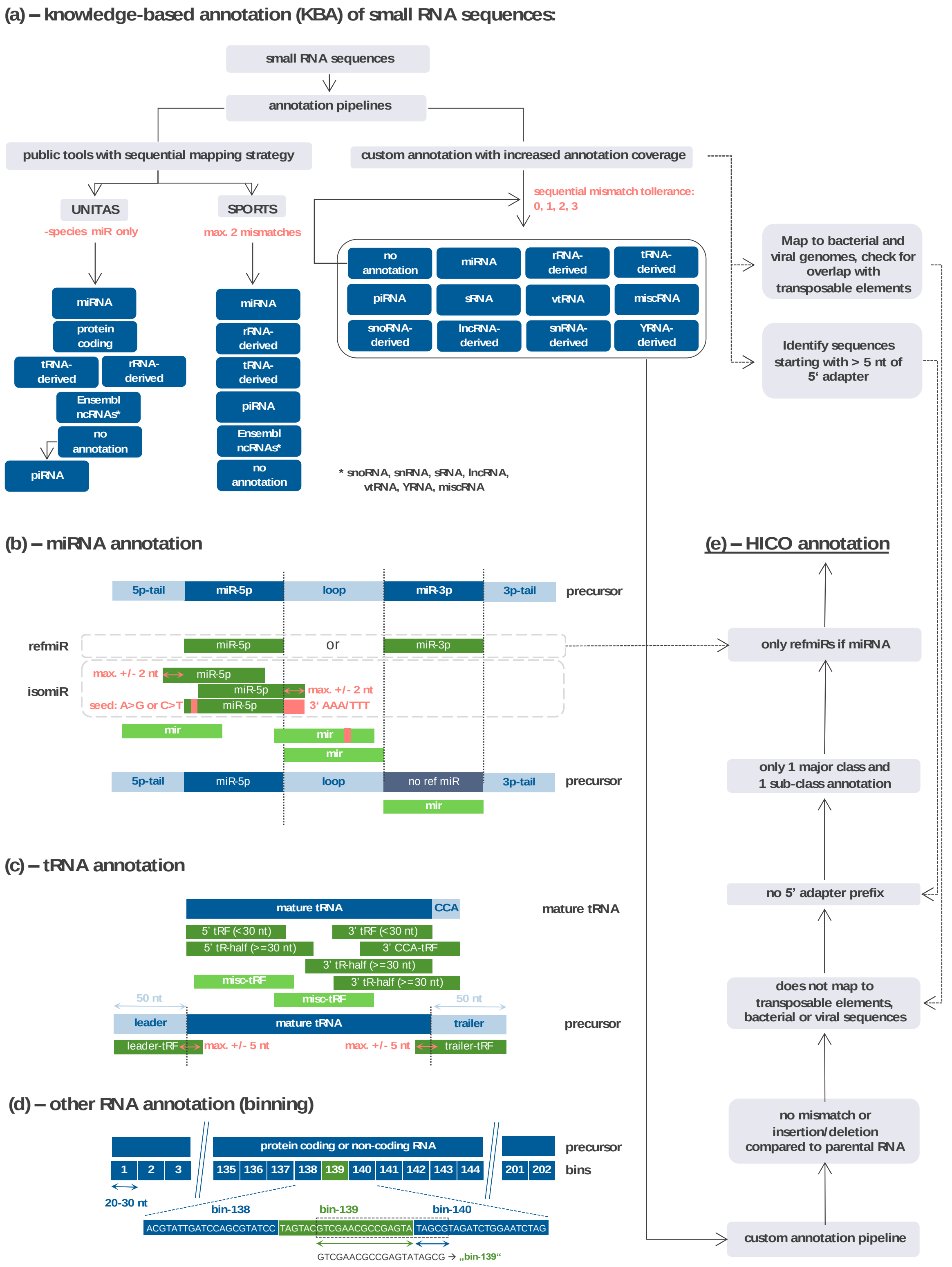


Figure S1. Overview of knowledge-based annotation strategy. (a) Small RNA sequences that were consistently detected in the healthy tissue samples of the TCGA miRNA-Seq data or the internal blood sample dataset were annotated using the published annotation tools SPORTS and UNITAS (left) and a custom annotation pipeline. UNITAS and SPORTS map the sequences sequentially to different small RNA class specific reference databases, which prioritizes the distinct small RNA classes and conceals potential assignment ambiguities. The custom annotation, in contrast, maps the sequences to the reference sequences of all small RNA classes at the same time starting with zero mismatch tolerance. Unmapped sequences are intended to map with iterating mismatch tolerance up to three mismatches. Additionally, all small RNA sequences are checked for potential bacterial or viral origin, for genomic overlap to human transposable element loci and whether they contain potential prefixes of the 5'-adapter. (b) Schematic overview of the miRNA annotation of the custom annotation (isomiR definition based on recent miRNA research). (c) Schematic overview of the tRNA annotation of the custom annotation (inspired by UNITAS sub-classification). (d) Binning strategy used in the custom annotation for the remaining RNA major classes. The number of nucleotides per bin is constant for each precursor sequence and ranges between 20 and 39 nucleotides. Assignments are based on the bin with the highest overlap to the sequence of interest. (e) Filtering steps that were applied to obtain the set of HICO annotations that were used for training of the TransfoRNA models.

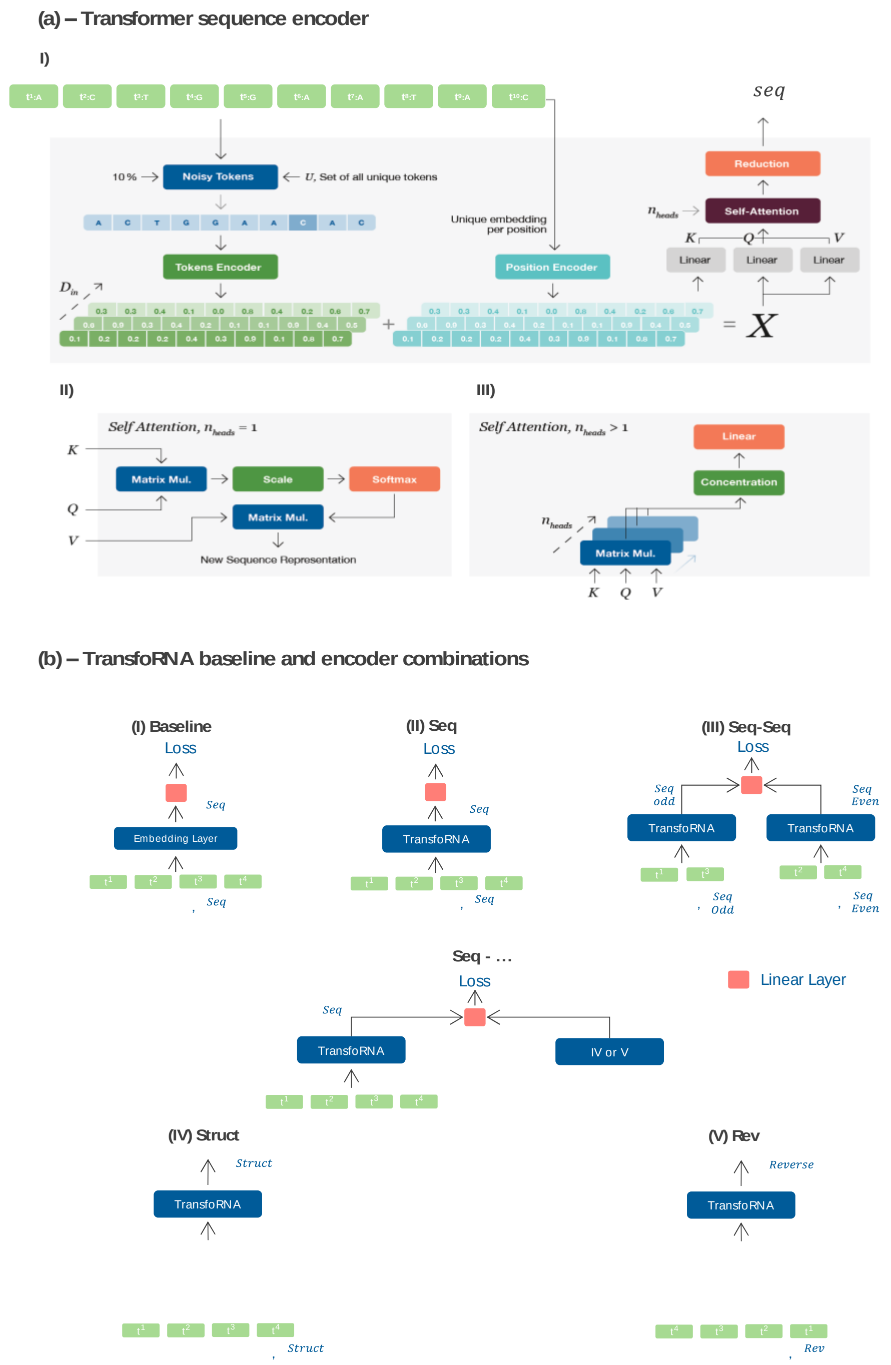


Figure S2. TransfoRNA architectural details. (a) An illustration of the TransfoRNA architecture. I) An example of a sequence (top left) is tokenized with window size of one before being passed to the transformer sequence encoder. A noise module is added to randomly replace 10% of the tokens with other tokens from the set of all unique tokens, U. A token encoder layer is then used to convert each token id to a vector space of dimension $D_{in}$. In parallel, the positional encoder encodes unique vectors of dimension $D_{in}$ per each unique position, independent of the input tokens. Both embeddings are then added and fed independently to three linear layers followed by a self-attention module which can be multi-headed if $n_{heads} > 1$. A reduction module is used to prevent the length factor from largely influencing sequence representations. (II) A depiction of inner function of the self-attention module that generates a new sequence representation while attending to all other tokens in the sequence. III) When $n_{heads}$>1, Multiheaded self-attention is used to compute a new sequence representation by concatenating each heads output followed by a linear layer.  (b) A schematic of all six models, each learning from a different input modality. Each input is converted to a reduced representation $R^{input}$, where the input is fed to a linear layer. Which outputs class predictions. The loss for all models is chosen to be the cross-entropy loss which computes the difference between the model predictions and the true classes (Major or sub-classes depending on the task). I) Baseline converts each token through an embedding layer to a continuous representation. All token embeddings are then concatenated and fed a linear layer. II) Seq uses the building block from (a) to convert the sequence into an embedding $R^{seq}$. III) For the Seq-Seq model, a sequence is tokenized and separated into odd-indexed and even-indexed token sets, where each set is then passed to a TransfoRNA block and converted to embeddings $R^{seq}_{even}$ and $R^{seq}_{odd}$. IV) Seq-Struct TransfoRNA takes as input the tokenized sequence and secondary structure (also presented as a sequence). Both sequences are converted to their reduced embedding representation $R^{seq}$ and $R^{struct}$ to which the loss is then applied. V) Seq-Rev takes as an input the tokenized sequence and its reverse.

**(a) – TransfoRNA performance on external long ncRNA benchmark dataset**

| model name | accuracy (%) | architecture | reference |
|---|---|---|---|
| nRC | 78.3 | CNN | (Fiannaca et al. 2017) |
| RNAGCN | 85.73 | GCN | (Rossi et al. 2019) |
| RPC-snRC | 95.38 | CNN | (Asim et al. 2021) |
| **Seq-Struct TransfoRNA** | **95.38** | **Transformer** | **this publication** |

**(b) –** Long ncRNA benchmark model (Seq-Struct TransfoRNA): performance on small RNAs (TCGA) is only 0.08 bACC.

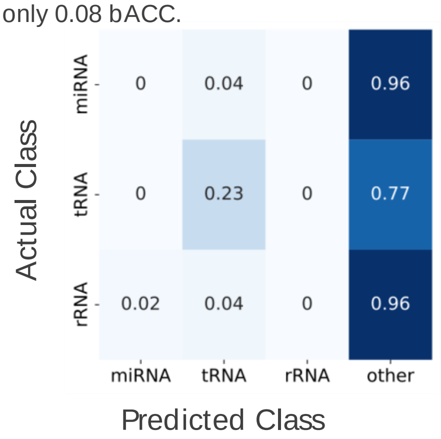


Figure S3. Evaluating the Transformer-based TransfoRNA model on a published benchmark of long non-coding RNA classification. (a) Competitive performance could be achieved with a sequence + structure model on the benchmark test set. (b) No predictive performance could be achieved when trying to annotate the TCGA small RNA data with the same model. The confusion matrix here is shown for the three major classes shared by the two datasets. CNN: Convolutional Neural Network; GCN: Graph Convolutional Network

**(a) – TransfoRNA performance (balanced accuracy) on HICO in-distribution (ID) test set**

| Model | Major class (bACC) | Sub class (bACC) | Major class, from sub-class (bACC) |
|---|---|---|---|
| Baseline | 52.83 ± 4.47 | 52.83 ± 1.01 | 89.61 ± 0.63 |
| Seq | 84.02 ± 3.65 | 97.70 ± 0.38 | 99.67 ± 0.14 |
| Seq-Seq | 77.36 ± 2.04 | 95.65 ± 0.50 | 99.41 ± 0.18 |
| Seq-Struct | 83.45 ± 4.05 | 97.71 ± 0.62 | 99.77 ± 0.07 |
| Seq-Rev | 83.51 ± 3.19 | 97.51 ± 0.34 | 99.71 ± 0.12 |

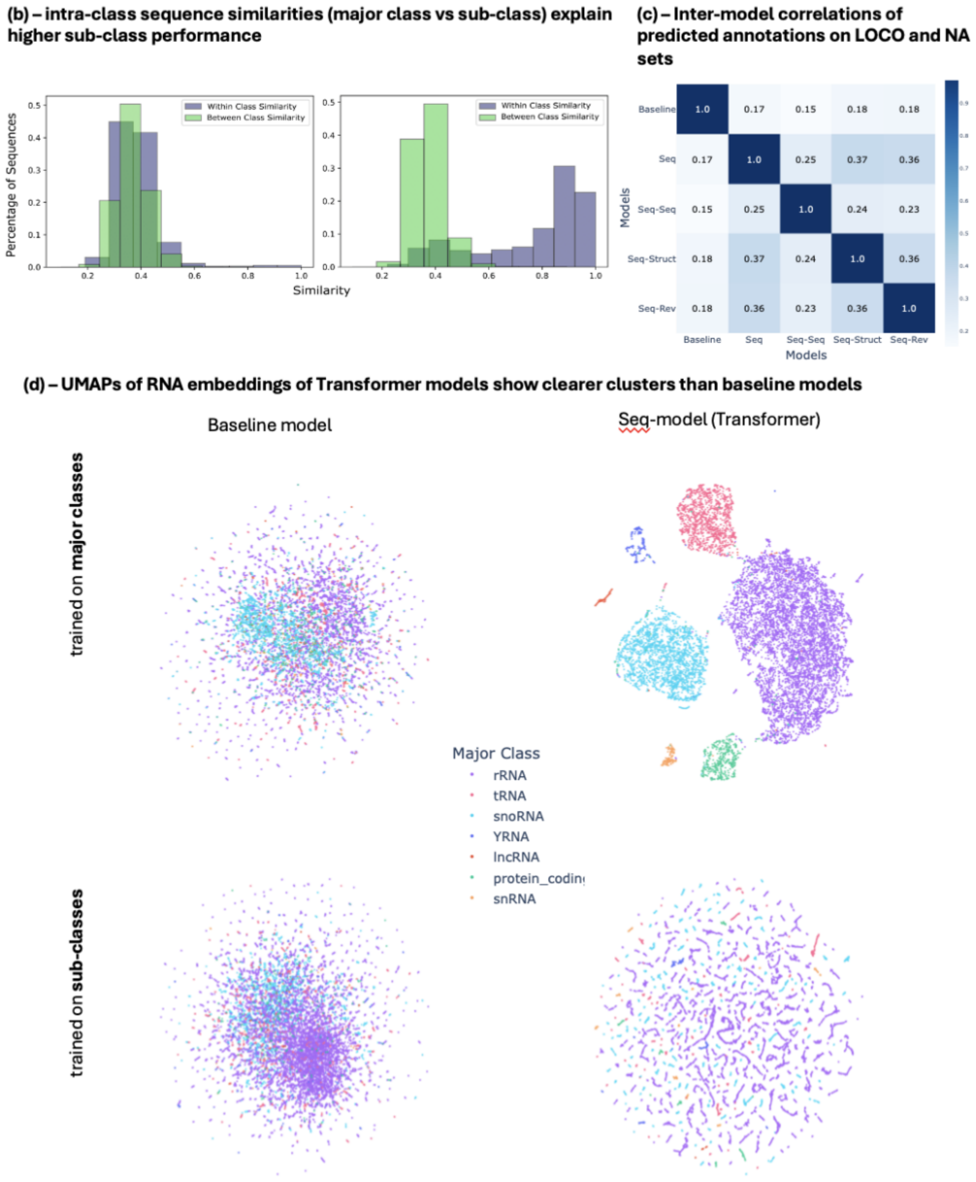

Figure S4. Results and analysis of TransfoRNA models on sub and major class classification: (a) Model performance using the in-distribution (ID) test set for models trained on the major-class and sub-class. (b) Two distributions of alignment-based sequence similarity scores, are shown per plot. “Within-class similarity” is the distribution of scores between all sequences sharing the same label, while “between-class similarity” reflects the distribution of similarity scores between all sets of sequences having different labels. Left: major class, Right: sub-class. (c) The correlation of sub-class predictions on the TCGA LOCO and NA sets for all models. (d) UMAPs of TransfoRNA embeddings of baseline and Seq models trained on the major classes (top) and sub-classes (bottom) belonging to 7 major classes. RNA sequences trained using the sub-classes are colored by major class as it is not feasible to show 374classes.

**(a) – different out-of-distribution (OOD) sets of RNAs, expected to be novel according to NLD**

| Set of novel RNAs (out-of-distribution, OOD) | category | Number of RNAs | Description |
|---|---|---|---|
| Rare sub-classes, including miRNAs | real RNA | 2919 | High-confidence annotations, but few (< 8) members per sub-class. Entire sub-classes held out from training. **Note**: miRNAs are only represented by a single annotated sequence per sub-class, and thus "rare" under this definition. |
| Putative 5'-adapter prefixes | real technical artefact | 294 | The 5'-end perfectly matches the last five or more nucleotides of the 5'-adapter sequence, commonly used in small RNA sequencing. |
| Recombined RNAs | computer-generated | 961 | RNA sequences from the training set were recombined into new sequences which retain a high sequence similiarity to members of the training set. |
| Random RNAs | computer-generated | 500 | Randomly chosen length and nucleotides with uniform probability. No overlap with the training set. |

**(b) – ID vs OOD NLD and AUC**

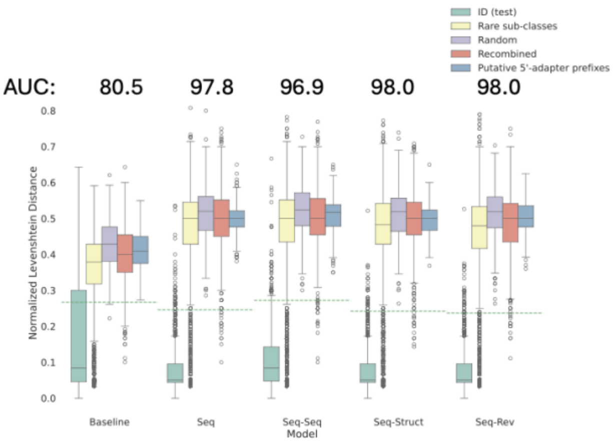


**(c) – Overlap between TransfoRNA annotations and low-confidence (LOCO) knowledge-based annotations**

| model | annotation overlap with LOCO set (%) | | |
|---|---|---|---|
| | for all RNAs | for RNAs familiar to TransfoRNA | for RNAs novel to TransfoRNA |
| Baseline | 8.60 | 60.86 | 0.00 |
| Seq | 10.37 | 78.88 | 0.90 |
| Seq-Seq | 9.97 | 74.21 | 0.47 |
| Seq-Struct | 9.99 | 76.60 | 0.81 |
| Seq-Rev | 10.22 | 76.88 | 0.76 |

Figure S5. Evaluation of the models trained on in-distribution (ID) sub-classes. (a) Definition of four different sets of RNAs that are known to be novel. (b) Distribution of the NLD, per model, and per split. The ID test set (sequences not used during training but belonging to familiar classes and therefore known to be familiar) along with four novel splits described in (a). The learned novelty threshold per model is depicted as a dashed horizontal line. (c) The novelty predictor was applied to the LOCO set of each model splitting them into two groups: familiar and novel. The percentage of overlap between TransfoRNA and LOCO annotations is given for all RNAs (familiar and novel), familiar RNAs only, and novel RNAs only.

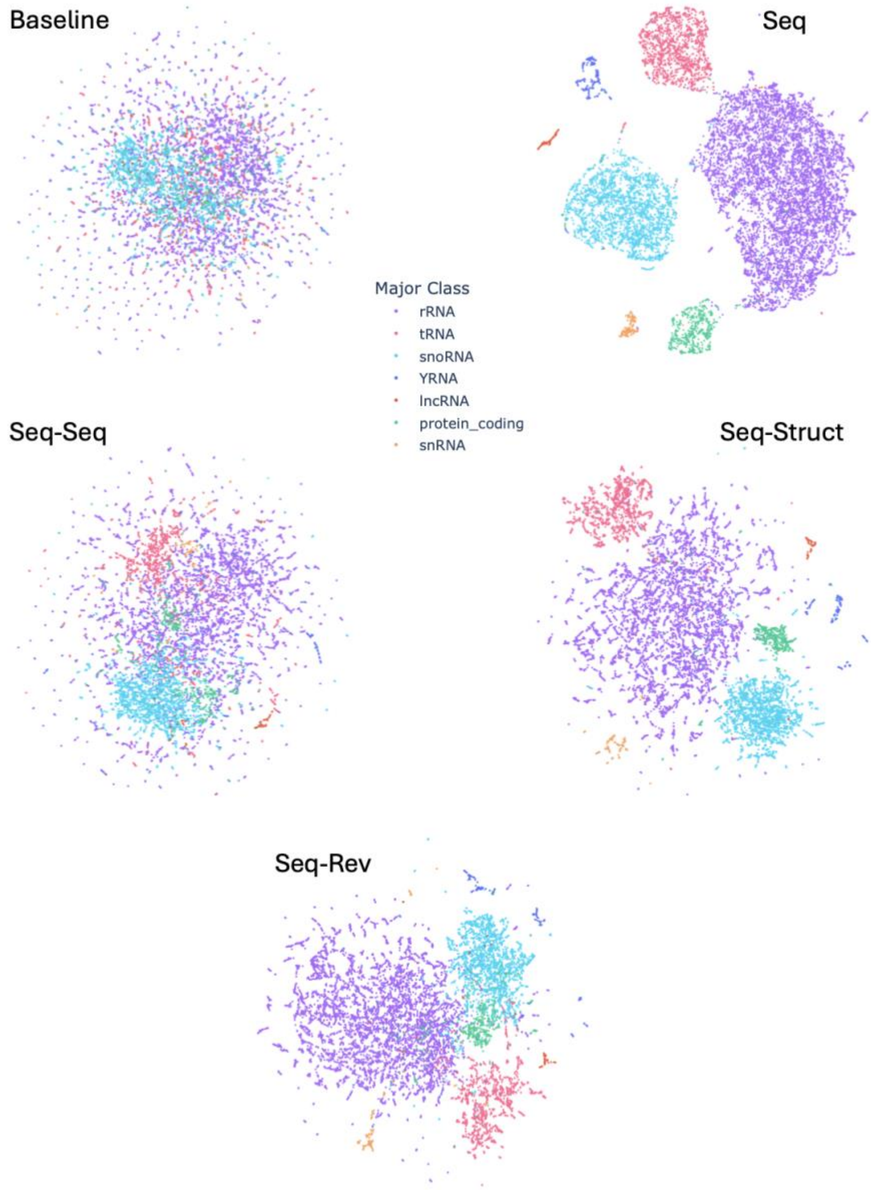


Figure S6. UMAPs of TransfoRNA embeddings of all models trained on high-confidence (HICO), in-distribution (ID), major-classes are shown.

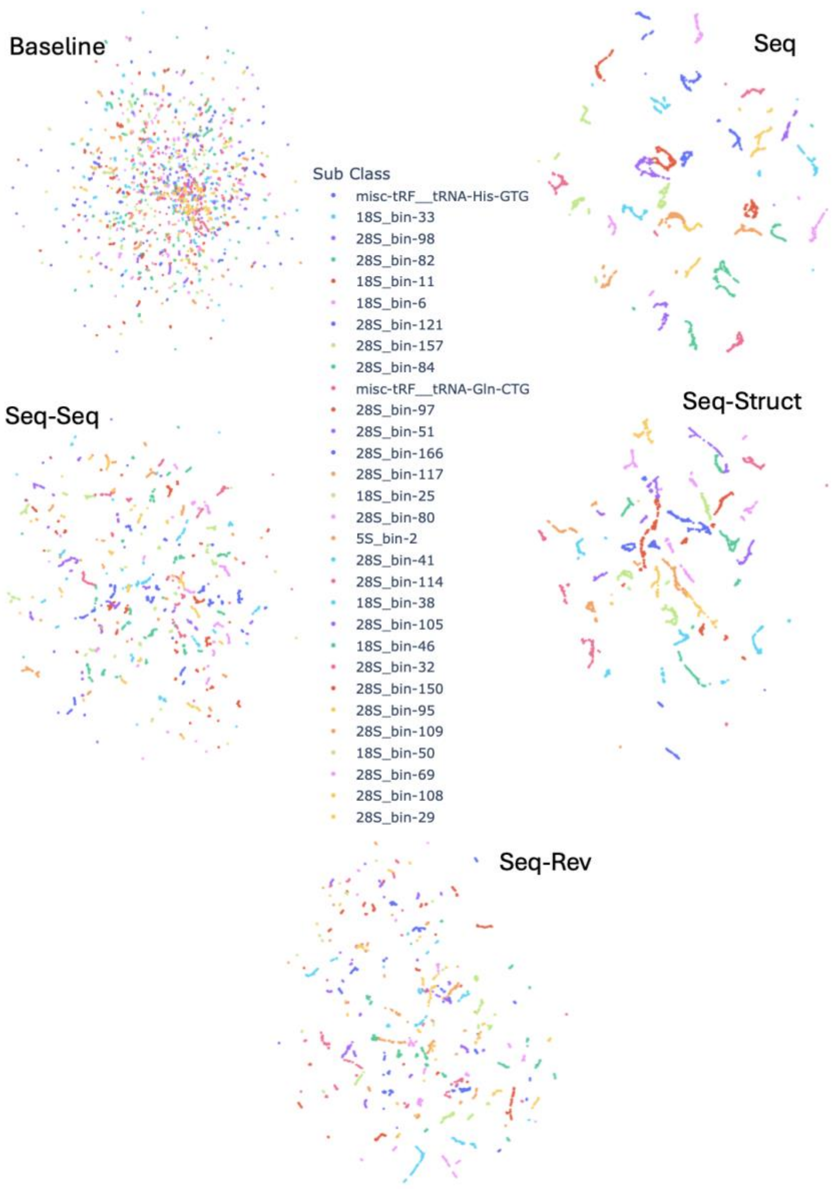


Figure S7. UMAPs of TransfoRNA embeddings of all models trained on high-confidence (HICO), in-distribution (ID), sub-classes are shown. Due to distinct color limitation, only samples belonging to the top 30 populated sub classes (out of 374) are shown.

1

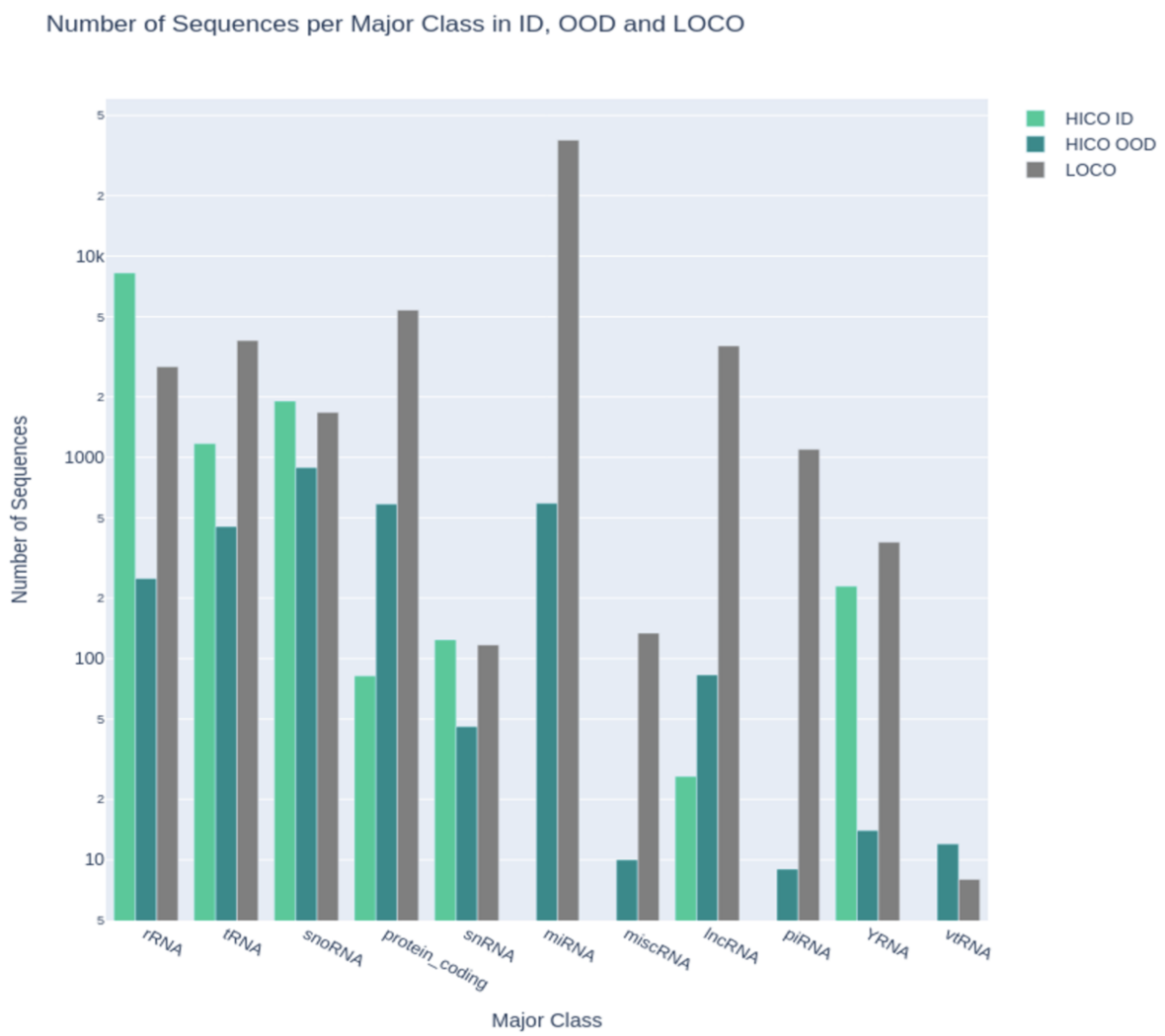


Figure S8. Number of sequences per major class. A depiction of all major classes used for sub-class classification task showing the proportion of sub-classes per each of the sets; high confidence in-distribution “HICO ID”, high confidence out-of-distribution ”HICO OOD” and the low confidence set “LOCO”.

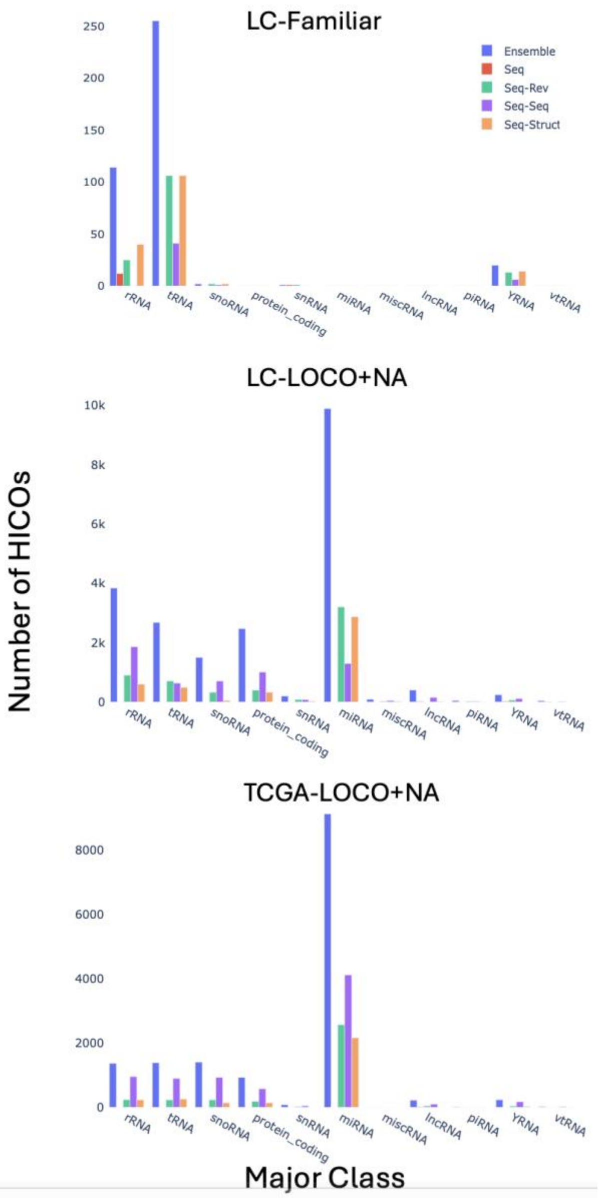


Figure S9. Advantage of Ensemble model over single models. For each of the three splits; a)LC-Familiar, b) LC_LOCO+NA and c) TCGA LOCO+NA, the number of hicos per major class per model is computed. Following that, the minimum number of hico sequences predicted by a given model per major class is deducted from the number of hico sequences predicted by other models, shown by a missing bar. Note: Baseline is not shown as Ensemble relies only on the Transformer-based models. The high jump in miRNAs could be seen in (b) and (c) but not in (a) because in LC-Familiar, only the canonical forms of miRNAs exist which means the miRNA sequences in LC-Familiar and in the training set (TCGA) were identical.

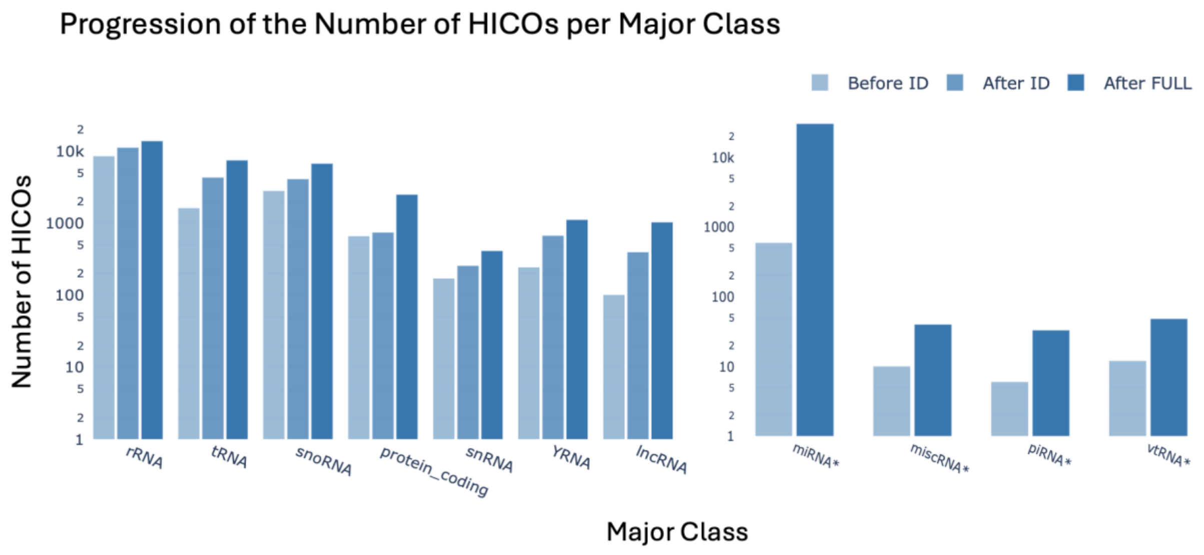


Figure S10. Progression of the number of high confidence sequences per major class. The first sevenmajor classes (left) were used during both phases, model development with held-out classes and training all HICO annotations in TCGA, while the rest of the major classes-marked with an asterisk-(right) were only used for training on TCGA.

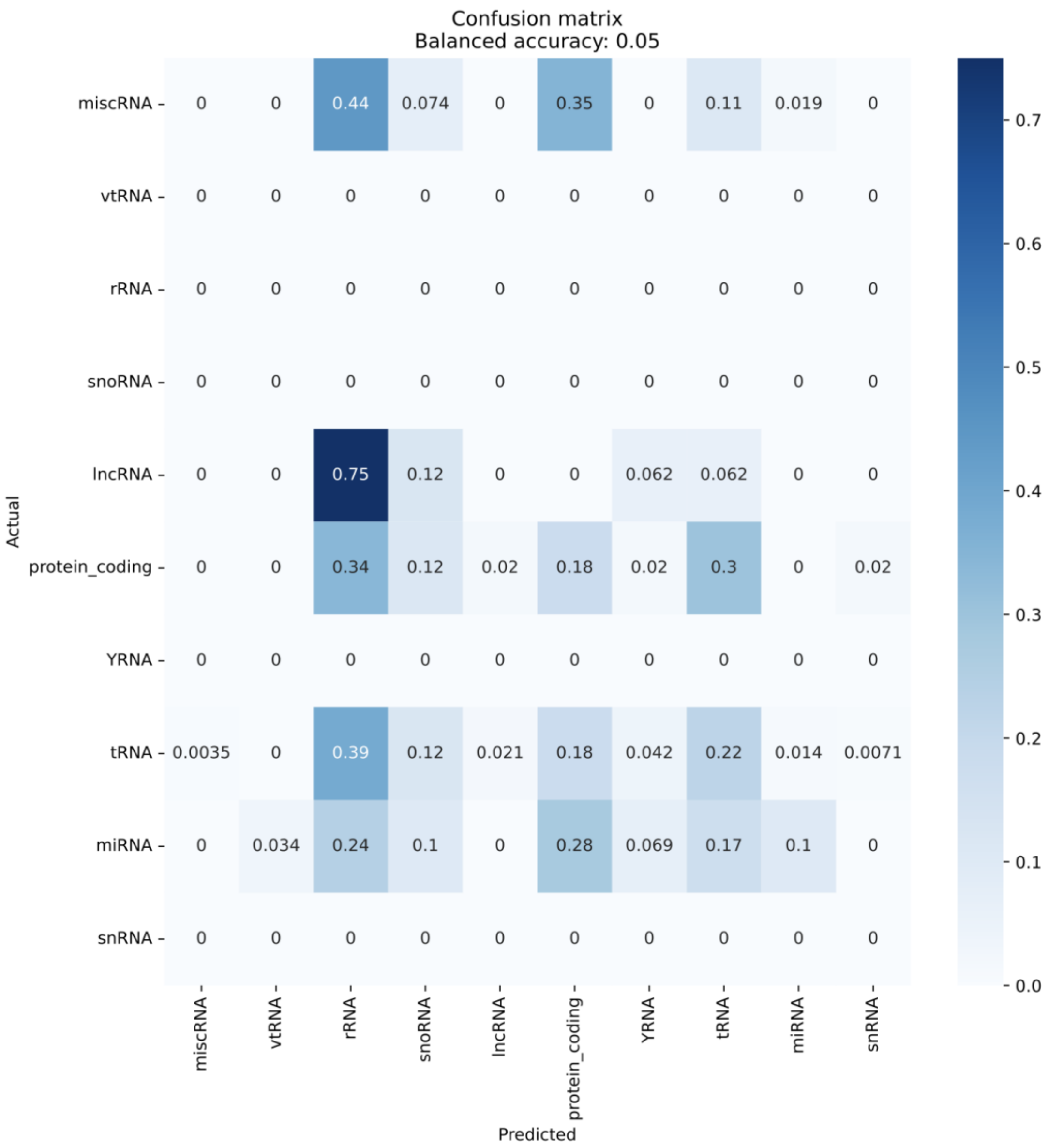


Figure S11. Confusion matrix of Ensemble model predictions on the LC-Novel set. Major class predictions are obtained by converting the models sub-class prediction to major class.

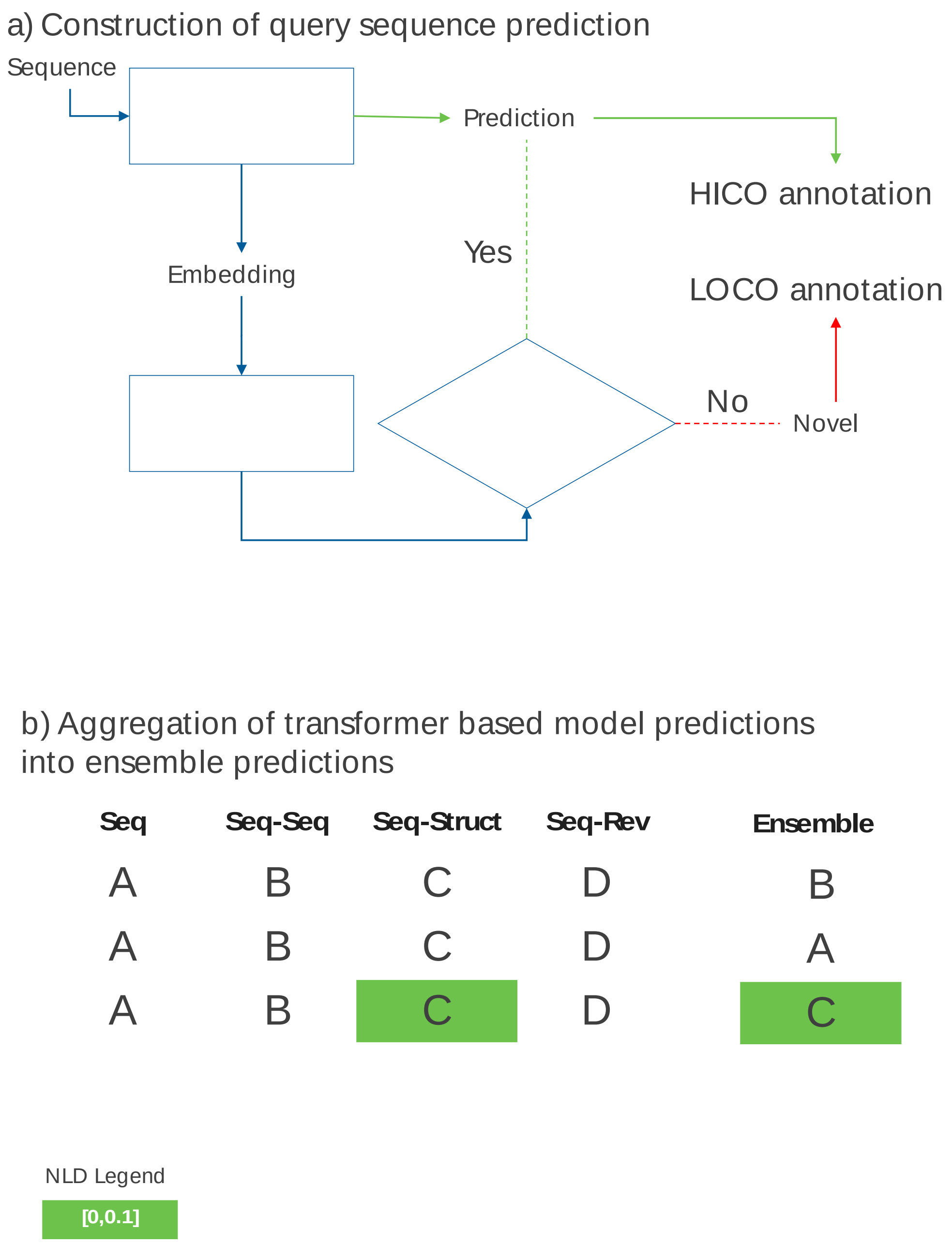


Figure S12. (a) Illustration of the proposed manner of fusing the TransfoRNA model predictions and the novelty prediction output. (b) Table shows different scenarios of how ensemble predictions are computed. A,B,C and D are the RNA example classes and the colors shows novelty predictors' output; Novel (red) and Familiar (green). The color intensity reflect the novelty predictors' confidence in its prediction computed based on the value of the NLD relative to learned threshold as shown in the legend.

Table S1. Thirteen different RNA classes present in the sncRNA benchmark dataset. Classes are listed along with the number of samples per class, as well as the maximum, median and minimum sequence length per class.

| **classes** | **No. of samples** | **max seq length** | **median seq length** | **min seq length** |
|---|---|---|---|---|
| IRES | 520 | 630 | 208 | 53 |
| Intron_gpI | 700 | 1182 | 320 | 133 |
| Leader | 700 | 237 | 127 | 38 |
| scaRNA | 700 | 445 | 139 | 78 |
| S5_rRNA | 700 | 199 | 119 | 61 |
| miRNA | 700 | 631 | 103 | 52 |
| tRNA | 700 | 177 | 73 | 47 |
| riboswitch | 700 | 399 | 132 | 44 |
| ribozyme | 700 | 1136 | 347 | 41 |
| S8_rRNA | 700 | 290 | 154 | 50 |
| CD-box | 700 | 404 | 96 | 54 |
| HACA-box | 700 | 508 | 133 | 59 |
| Intron_gpII | 700 | 241 | 90 | 48 |

Table S2. An ablation study illustrating the importance of different components of the Seq-Struct TransfoRNA model. Models were trained in a supervised manner on the sncRNA dataset. Each highlighted block shows the effect of changing the values of a given hyperparameter shown in the first row. Token lengths were varied from 1 to 5 while the number of heads, $n_{heads}$, of the multi-headed attention was set to either 1 or 4. The effect of the false input module was also tested. The variance in loss was the highest with varying token length followed by changing the loss scheme. The highest accuracy score was achieved with token length = 4, $n_{heads}$ = 4, using only the contrastive loss and the incorporation of the false input module.

| | **token length** | | | | | **n-heads** | | **false input** | | **accuracy(%)** |
|---|---|---|---|---|---|---|---|---|---|---|
| | 1 | 2 | 3 | 4 | 5 | 1 | 4 | No | Yes | |
| Yes | | | | | | Yes | Yes | | 90.15 | |
| | Yes | | | | | Yes | Yes | | 91.96 | |
| | | Yes | | | | Yes | Yes | | 91.96 | |
| | | | Yes | | | Yes | Yes | | 94.42 | |
| | | | | Yes | | Yes | Yes | | 94.23 | |
| | | | Yes | | Yes | | Yes | | 94.02 | |
| | | | Yes | | | Yes | Yes | | 94.42 | |
| | | | Yes | | | Yes | Yes | | 94.39 | |
| | | | Yes | | | Yes | Yes | | 95.33 | |
| | | | Yes | | | Yes | Yes | | 95.33 | |
| | | | Yes | | | Yes | | Yes | 95.42 | |

Table S4: RNA count with putative 5′-adapter affixes of length > 4. TCGA: The Cancer Genome Atlas, LC: lung cancer.

| **5’-adapter prefix** | **Match length** | **Count in dataset A (TCGA)** | **Count in dataset B (LC)** |
|---:|---|---|---|
| CGATC | 5 | 25 | 260 |
| ACGATC | 6 | 1 | 84 |
| GACGATC | 7 | 0 | 207 |
| CGACGATC | 8 | 0 | 136 |
| CCGACGATC | 9 | 0 | 73 |
| TCCGACGATC | 10 | 1 | 47 |
| GTCCGACGATC | 11 | 8 | 358 |
| AGTCCGACGATC | 12 | 26 | 332 |
| CAGTCCGACGATC | 13 | 34 | 219 |
| ACAGTCCGACGATC | 14 | 27 | 243 |
| TACAGTCCGACGATC | 15 | 26 | 196 |
| CTACAGTCCGACGATC | 16 | 32 | 274 |
| TCTACAGTCCGACGATC | 17 | 33 | 238 |
| TTCTACAGTCCGACGATC | 18 | 28 | 140 |
